# Developmental genetics of corolla tube formation: role of the tasiRNA-ARF pathway and a conceptual model

**DOI:** 10.1101/253112

**Authors:** Baoqing Ding, Rui Xia, Qiaoshan Lin, Vandana Gurung, Janelle M. Sagawa, Lauren E. Stanley, Matthew Strobel, Pamela K. Diggle, Blake C. Meyers, Yao-Wu Yuan

## Abstract

More than 80,000 angiosperm species produce flowers with petals fused into a corolla tube. As an important element of the tremendous diversity of flower morphology, the corolla tube plays a critical role in many specialized interactions between plants and animal pollinators (e.g., beeflies, hawkmoths, hummingbirds, nectar bats), which in turn drives rapid plant speciation. Despite its clear significance in plant reproduction and evolution, the corolla tube remains one of the least understood plant structures from a developmental genetics perspective. Through mutant analyses and transgenic experiments, here we show that the tasiRNA-ARF pathway is required for corolla tube formation in the monkeyflower species *Mimulus lewisii*. Loss-of-function mutations in the *M. lewisii* orthologs of *ARGONAUTE7* and *SUPPRESSOR OF GENE SILENCING 3* cause a dramatic decrease in abundance of *TAS3*-derived small RNAs and a moderate up-regulation of *AUXIN RESPONSE FACTOR 3* (*ARF3*) and *ARF4*, which lead to inhibition of lateral expansion of the bases of petal primordia and complete arrest of the upward growth of the inter-primordial regions, resulting in unfused corollas. By using an auxin reporter construct, we discovered that auxin distribution is continuous along the petal primordium base and the inter-primordial region during the critical stage of corolla tube formation in the wild-type, and that this auxin distribution is much weaker and more restricted in the mutant. Together, these results suggest a new conceptual model highlighting the central role of auxin directed synchronized growth of the petal primordium base and the inter-primordial region in corolla tube formation.

## INTRODUCTION

About one third of the ~275,000 angiosperm species produce flowers with petals fused into a corolla tube (i.e., sympetalous), forming a protective enclosure of nectaries and reproductive organs. Corolla tubes have evolved multiple times independently across the angiosperm tree of life (Endress, 2011), most notably in the common ancestor of the Asterids, a clade containing more than 80,000 species (Schonenberger and Von Balthazar, 2013). Subsequent elaboration in length, width, and curvature has led to a great variety of corolla tube shapes that enabled asterid species to exploit many specialized pollinator groups (e.g., beeflies, hawkmoths, hummingbirds, nectar bats), which in turn drives rapid plant speciation (Muchhala, 2006; Hermann and Kuhlemeier, 2011; Paudel et al., 2015; Lagomarsino et al., 2016). As such, the corolla tube has long been considered a key morphological innovation that contributed to the radiation of the Asterids (Endress, 2011). Despite its critical importance in the reproductive success and evolution of such a large number of species, the corolla tube remains one of the least understood plant structures from a developmental genetics perspective (Specht and Howarth, 2015; Zhong and Preston, 2015).

Historically, corolla tube formation has been the subject of extensive morphological and anatomical studies (Boke, 1948; Kaplan, 1968; Govil, 1972; Nishino, 1976, 1978, 1983a, 1983b; Erbar, 1991; Erbar and Leins, 1996; Kajita and Nishino, 2009; El Ottra et al., 2013). In particular, numerous studies have described the detailed ontogenetic process of corolla tube development in one subgroup of the asterid clade, the Lamiids, which contains some classical plant genetic model systems such as snapdragon (*Antirrhinum*), petunia (*Petunia*), and morning glory (*Ipomoea*) (Govil, 1972; Nishino, 1976, 1978, 1983a, 1983b; Singh and Jain, 1979; Erbar, 1991; Vincent and Coen, 2004; Kajita and Nishino, 2009; Erbar and Leins, 2011). A common theme emerging from these studies is that during the early stage of petal development, petal primordia are initiated separately, followed by rapid extension of the petal bases toward the inter-primordial regions, which also grow coordinately, causing congenital “fusion” of the petal primordia and formation of the corolla tube. Little is known, however, about the genetic control of this early-phase lateral extension of the petal base or the nature of the coordinated inter-primordial growth.

To date only a few genes have been implicated in corolla tube formation. Loss-of-function alleles of the *FEATHERED* gene in Japanese morning glory (*Ipomoea nil*) and the *MAEWEST* gene in petunia (*Petunia × hybrida*), both generated by transposon insertions, result in unfused corollas (Iwasaki and Nitasaka, 2006; Vandenbussche et al., 2009). *FEATHERED* and *MAEWEST* encode KANADI and WOX transcription factors, and their *Arabidopsis* orthologs are *KANADI1* and *WOX1*, respectively. In addition, ectopic expression of the *Arabidopsis* TCP5 protein fused with a repressor motif in *Ipomoea* also disrupted corolla tube formation (Ono et al., 2012). However, whether the endogenous *TCP5* ortholog in *Ipomoea* is involved in corolla tube development is unclear. More recently, it was reported that transient knock-down of the *Petunia* NAC-transcription factors *NAM* and *NH16* via virus-induced gene silencing (VIGS) also caused decreased petal fusion (Zhong et al., 2016), but the interpretation of this result was confounded by the observation that occasional flowers produced on the “escape shoots” of the loss-of-function *nam* mutants have normal corolla tubes (Souer et al., 1996). The fact that these genes were characterized from different plant systems and through different methods (transposon insertion alleles, heterologous expression of chimeric repressor, and VIGS) makes it challenging to interpret their genetic relationships and their precise functional roles in corolla tube formation.

One way to overcome this problem is to systematically analyze corolla tube mutants in a single model system. To this end, we have employed a new genetic model system, the monkeyflower species *Mimulus lewisii*, mainly for its ease in chemical mutagenesis and *Agrobacterium-mediated in planta* transformation (Owen and Bradshaw, 2011; Yuan et al., 2013a). *M. lewisii* is a typical bumblebee-pollinated species with a conspicuous corolla tube (Figure 1A). Through ethyl methanesulfonate (EMS) mutagenesis, we have generated a dozen recessive mutants (named *flayed*) with split corolla tubes. Here we report the characterization of one group of mutants, caused by loss-of-function mutations in two genes that are required for the biogenesis of trans-acting short interfering RNAs (tasiRNAs).

**Figure 1.**
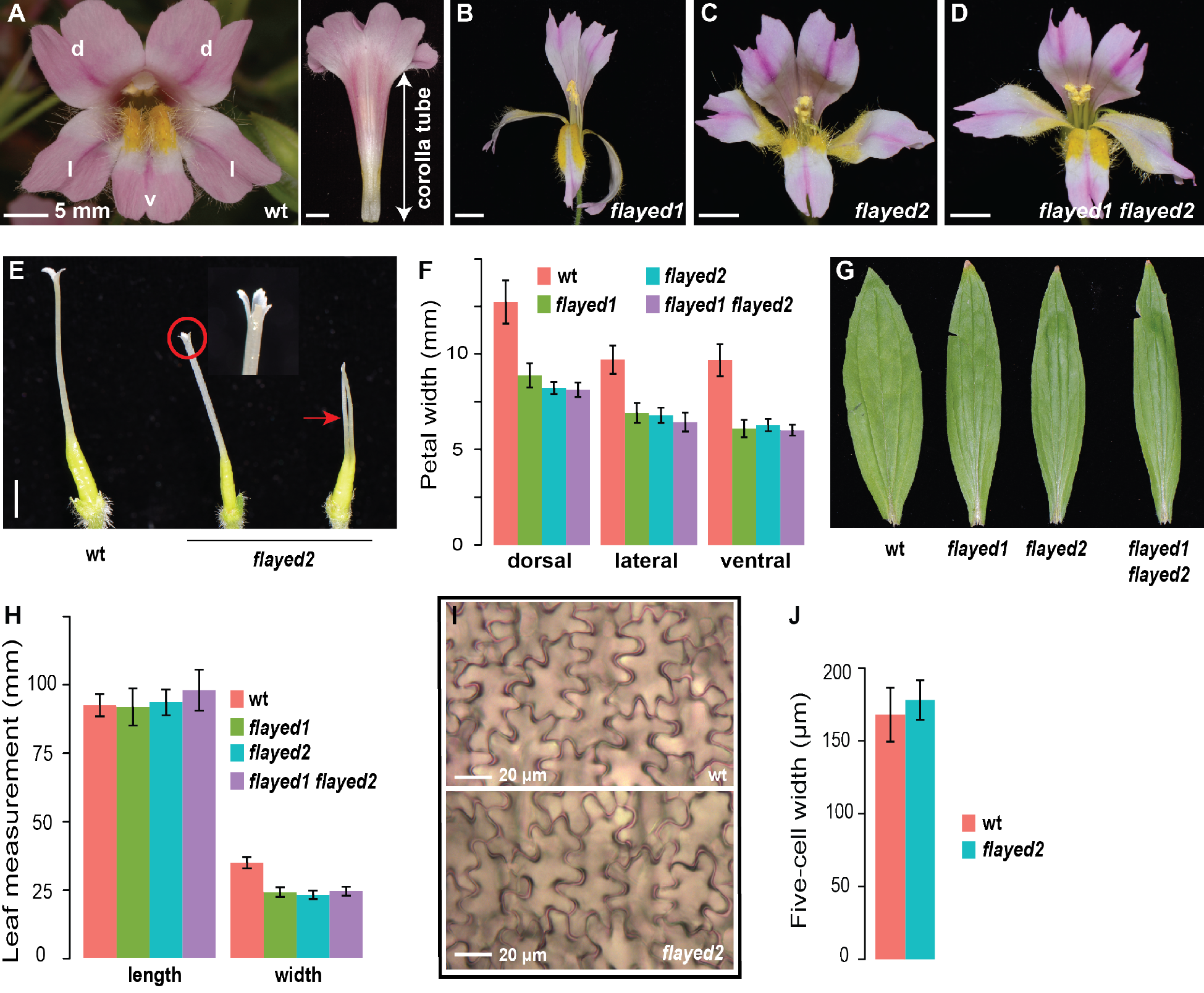
Phenotypic characterization of the *flayed* mutants. (**A**) Face and side view of the wild-type *M. lewisii* (inbred line LF10) corolla. d: dorsal; l: lateral; v: ventral. (**B**) Side view of *flayed1* corolla. (**C** and **D**) Face view of *flayed2* (**C**) and the double mutant (**D**). (**E**) Pistil of the wild-type (wt) and *flayed2*. The pistil phenotype of *flayed1* and the double mutant is the same as that of *flayed2*. (**F**) Quantitative comparison of petal width in wt (n =18), *flayed1* (n = 10), *flayed2* (n = 12), and the double mutant (n = 12). Detailed measurement data are presented in Table S1. (**G**) Overall shape of the fourth leaf (the largest leaf) of mature plants. (**H**) Quantitative comparison of length and width of the fourth leaf, with the same sample sizes as in (**F**). (**I**) Abaxial epidermal cells of dorsal petal lobes. (**J**) Width of five contiguous abaxial epidermal cells of the dorsal petal lobes in the wt (n = 15) and *flayed2* (n = 15). Scale bars in (**A-D**) are 5 mm. Error bars in (**F**), (**H**), and (**J**) are 1 SD.

Among the tasiRNA loci characterized to date, *TAS3* is the most widely conserved, found in virtually all land plants (Xia et al., 2017). *TAS3* transcript bears two binding sites for miR390, which triggers the production of phased tasiRNAs, including the highly conserved “tasiARF” that targets *AUXIN RESPONSE FACTOR 3* (*ARF3*) and *ARF4* (Allen et al., 2005; Axtell et al., 2006). This tasiRNA-ARF regulatory module has been shown to play a critical role in leaf adaxial/abaxial polarity and blade expansion (i.e., lamina growth) in both eudicots (Fahlgren et al., 2006; Garcia et al., 2006; Hunter et al., 2006; Yan et al., 2010; Yifhar et al., 2012; Zhou et al., 2013) and monocots (Nagasaki et al., 2007; Nogueira et al., 2007; Douglas et al., 2010). Consistent with previous studies, here we demonstrate that in the *M. lewisii* mutants, *TAS3*-derived tasiRNAs decrease dramatically in abundance and *MlARF3* and *MlARF4* expression are upregulated. Importantly, we show that malfunction of the tasiRNA-ARF pathway in the *M. lewisii* mutants impedes the early lateral expansion of the petal primordium bases and the coordinated inter-primordial growth, most likely through change of auxin homeostasis, leading to unfused petal primordia. Integrating our molecular and phenotypic analyses of the tasiRNA-ARF pathway and auxin accumulation patterns in *M. lewisii* with historical insights from morphological and anatomical studies of various sympetalous species, we propose a new conceptual model for the genetic control of corolla tube formation, which offers logical connections among the sporadic previous reports of corolla tube mutants and makes clear predictions that can be readily tested using the *Mimulus* system.

## RESULTS

### Phenotypic Characterization of the *flayed1* and *flayed2* Mutants

Three of the recessive mutants recovered from EMS mutagenesis using the inbred line LF10, *flayed1-flayed3*, are morphologically indistinguishable. Pair-wise crosses suggested that they belong to two complementation groups, *flayed1* and *flayed2* (*flayed3* is allelic to *flayed2*) (Figure 1B and C). In addition to having unfused petals, these mutants display carpel fusion defects, with phenotypes varying from flower to flower within the same plant. Most mutant flowers have two fused carpels, as in the wild-type, but have partially split stigmas with more than two lobes (Figure 1E). Less frequently there are flowers with two almost completely separate styles. The length of mutant pistils is also reduced compared to the wild-type (Figure 1E). No obvious phenotypes were observed in the stamens of these mutants.

Another notable feature of the *flayed1/2* mutants is the reduced width of lateral organs. The dorsal and lateral petals show ~30% decrease in width compared to the wild-type, and the ventral petal shows ~37% decrease (Figure 1F; Table S1). Leaf width is also substantially reduced (by ~40%) in the mutants, but length is unaffected (Figure 1G and H; Table S1). To determine whether the reduction in petal width is due to change in cell number, cell size, or both, we measured the width of abaxial epidermal cells of the dorsal petal lobe for both the wild-type and the *flayed2* mutant. Because the petal lobe abaxial epidermal cells are irregularly shaped (Figure 1I), the width measurements were done on five contiguous cells to account for the variation among individual cells within the same sample. No significant difference in cell width was found between the wild-type and *flayed2* (Figure 1J), which suggests that the difference in petal width between the mutant and the wild-type is primarily due to difference in cell number (i.e., number of cell divisions).

Unlike the morning glory mutant *feathered* (Iwasaki and Nitasaka, 2006) or the petunia mutant *maewest* (Vandenbussche et al., 2009), *flayed1/2* do not show any defects in tissue adaxial/abaxial polarity. Instead, the *flayed1/2* mutants closely resemble the petunia mutant *choripetala suzaane* (*chsu*), which also have split corolla tubes, variable carpel fusion defects, and narrower leaf with normal adaxial/abaxial polarity. Unfortunately, the molecular identity of *CHSU* is still unknown.

### *FLAYED1* and *FLAYED2* Are the Orthologs of Arabidopsis *AGO7*and *SGS3*, Respectively

To identify the causal genes of *flayed1* and *flayed2*, we analyzed each mutant using a genomic approach that combines the advantages of bulk segregant analysis and comparison of single nucleotide polymorphism (SNP) profiles between multiple EMS mutants (**Methods**), as demonstrated in a previous study (LaFountain et al., 2017). We narrowed the causal mutation of *flayed1* and *flayed2* down to 38 and 19 candidate SNPs, respectively (Table S2 and S3). The vast majority of these SNPs locate in non-coding, repetitive sequences, with only two or three mutations resulting in amino acid changes in each mutant (Table S2 and S3). Notably, in both *flayed1* and *flayed2*, there is one mutation leading to a premature stop codon, in the ortholog of Arabidopsis *ARGONAUTE7* (*AGO7*) and *SUPPRESSOR OF GENE SILENCING 3* (*SGS3*), respectively (Figure 2A and B; Table S2 and S3). *AGO7* and *SGS3* are part of the same tasiRNA biogenesis pathway (Peragine et al., 2004; Yoshikawa et al., 2005; Chen, 2010), which would explain the indistinguishable mutant phenotypes of *flayed1* and *flayed2*. Furthermore, sequencing the coding DNA (CDS) of *MlSGS3* in *flayed3*, which is allelic to *flayed2*, revealed an independent mutation that also leads to a premature stop codon (Figure 2B). Together, these results suggested that *MlAGO7* and *MlSGS3* were the most promising candidate genes for *FLAYED1* and *FLAYED2*, respectively.

**Figure 2.**
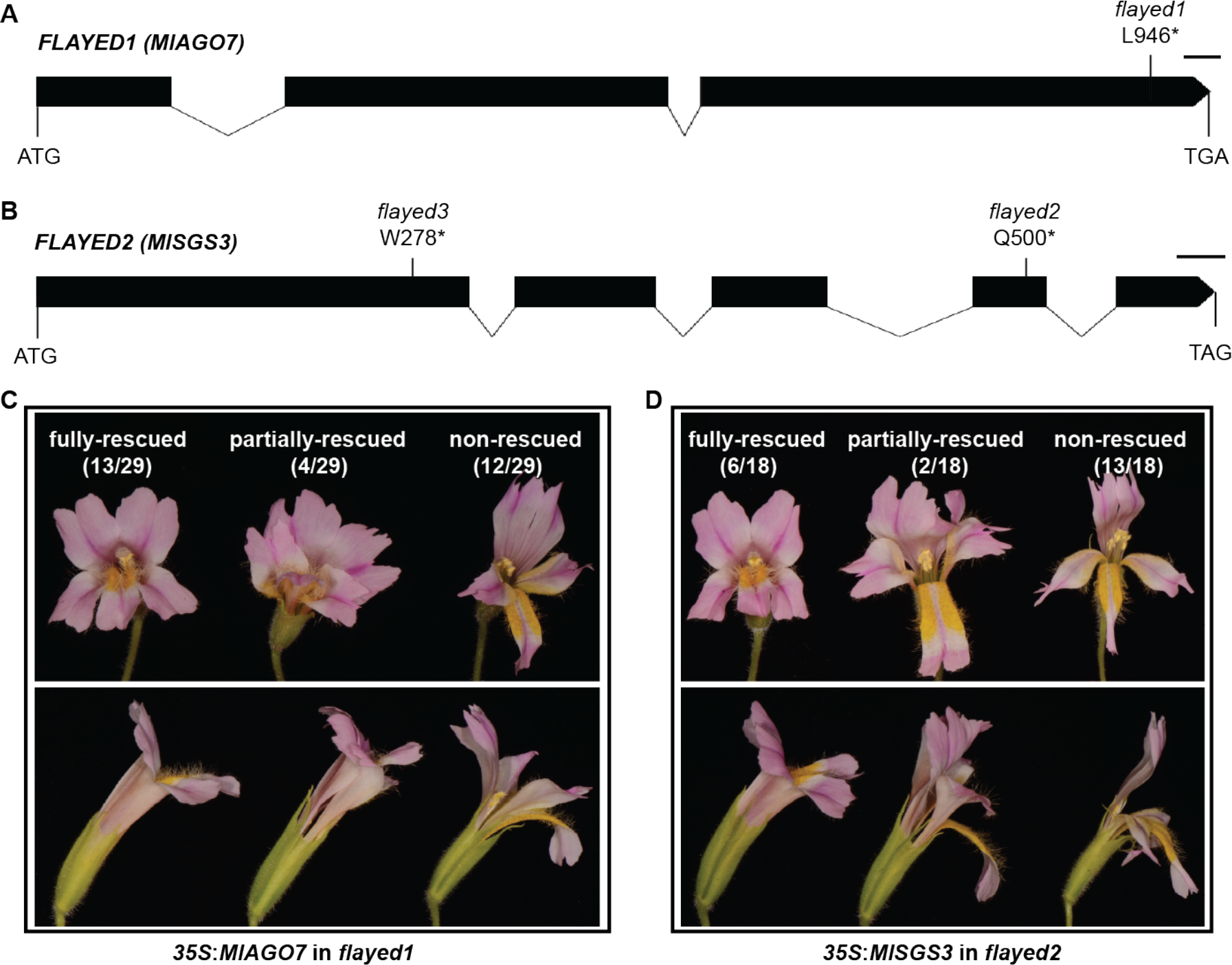
*FLAYED1* and *FLAYED2* encode the orthologs of *Arabidopsis* AGO7 and SGS3, respectively. (**A** and **B**) Schematics of *MlAGO7* (**A**) and *MlSGS3* (**B**) gene structure, with causal mutations indicated. Black box: coding DNA; Line: Intron. Scale bars are 100 bp. (**C** and **D**) Flower phenotypes of *35S:MlAGO7* (**C**) and *35S:MlSGS3* (**D**) transgenics in the *flayed1* and *flayed2* mutant background, respectively (top: face view; bottom: side view). The proportion of fully-rescued, partially-rescued, and non-rescued lines are shown in the parentheses.

To verify gene identities, full-length CDS of *MlAGO7* and *MlSGS3* were introduced to the *flayed1* and *flayed2* mutant background, respectively, driven by the cauliflower mosaic virus 35S promoter. Among the 29 independent *35S:MlAGO7* lines in the *flayed1* background, 13 showed a fully rescued phenotype that is indistinguishable from the wild-type; four lines showed a partially rescued phenotype, with petal and leaf width indistinguishable from wild-type but the petals remained unfused (Figure 2C). Similarly, six of the 18 *MlSGS3* over-expression lines in the *flayed2* background displayed a fully rescued phenotype and two displayed a partially rescued phenotype (Figure 2D). qRT-PCR assessment of *MlAGO7* and *MlSGS3* expression in 5mm floral buds showed that, in the fully rescued lines, expression levels of the transgenes are 4~64-fold higher than those of the corresponding endogenous genes (Figure S1). These results confirmed that *MlAGO7* and *MlSGS3* are indeed the causal genes underlying *flayed1* and *flayed2*, respectively.

Knowing the causal genes and mutations allowed direct genotyping of a “*flayed1* × *flayed2*” F_2_ population to identify *flayed1 flayed2* double mutants, which are phenotypically indistinguishable from the single mutants (Figure 1D, F, G, H). This further indicates that *MlAGO7* and *MlSGS3* function in the same genetic pathway in *Mimulus*, as expected.

### The *flayed1/2* Phenotypes Are Primarily Mediated Through the tasiRNA-ARF Pathway

Because *AGO7* and *SGS3* are necessary components of the miR390-TAS3-ARF pathway (Figure 3A), and the highly conserved, *TAS3*-derived tasiARFs are known to play a critical role in leaf polarity and lamina growth by repressing *ARF3/4* expression, we hypothesized that the *flayed1/2* phenotypes (e.g., reduced width of lateral organs) are primarily mediated through the tasiRNA-ARF pathway. This hypothesis makes three clear predictions: (**i**) The abundance of *TAS3*-derived small RNAs, including tasiARFs, should be much lower in the mutants compared to the wild-type; (**ii**) The *M. lewisii* orthologs of *ARF3/4* should be upregulated in the mutants; and, (**iii**) Artificial upregulation of the *M. lewisii ARF3/4* orthologs in the wild-type background should recapitulate the *flayed1/2* phenotypes.

**Figure 3.**
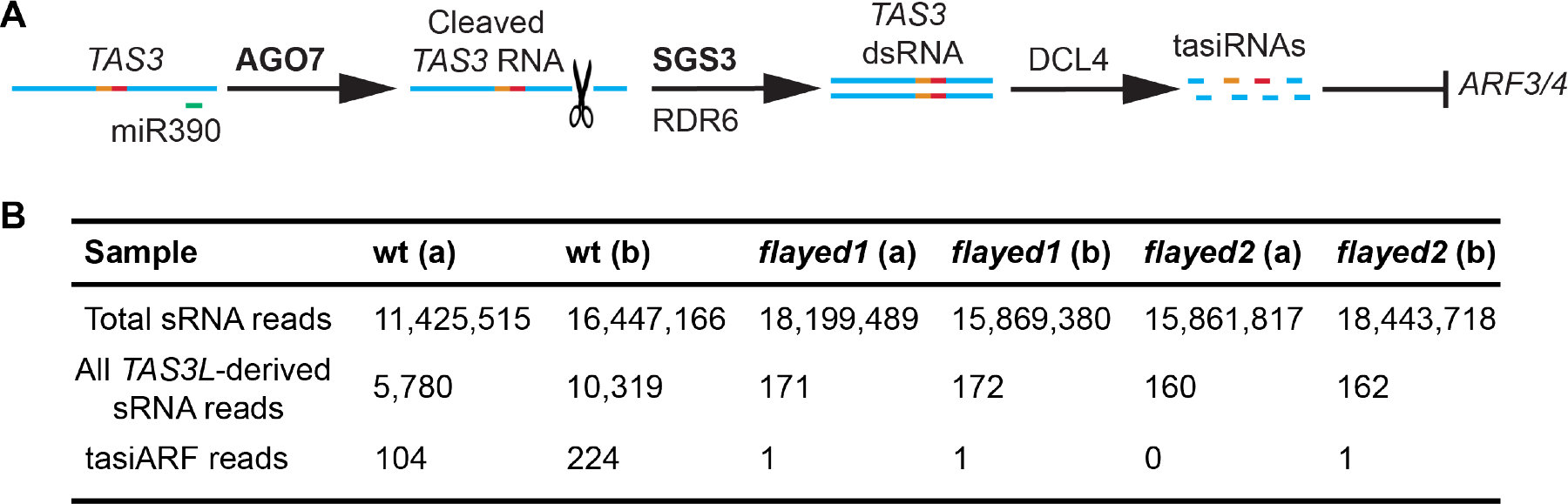
Small RNA analysis. (**A**) Schematic of the *miR390-TAS3-ARF* pathway. The orange and red lines represent the two tandem tasiARFs (see Figure S2 for detailed annotations). **(B)**Small RNA counts in the wild-type (wt), *flayed1*, and *flayed2*. Two biological replicates were sequenced for each genotype.

To test the first prediction, we sequenced the total small RNA pool from young floral buds (5-mm) of the wild-type, *flayed1*, and *flayed 2*. Like most other angiosperms, *M. lewisii* has two kinds of *TAS3* genes (each represented by only a single copy in the *M. lewisii* genome): *TAS3S* contains a single, centrally located tasiARF, whereas *TAS3L* contains two tandem tasiARFs (Xia et al., 2017) (Figure S2). No *TAS3S*-derived small RNAs were detected in any of the sequenced samples, suggesting that the *TAS3S* gene is not expressed. *TAS3L*-derived small RNAs were detected at the level of ~600 per million reads in the wild-type, but decreased >50-fold in both *flayed1* and *flayed2* (Figure 3B). In particular, the tasiARFs were almost entirely absent from the mutant samples (Figure 3B). These results confirmed the first prediction.

To test the second prediction, we first searched the *M. lewisii* genome for *ARF3/4* homologs and found a single ortholog for each of the two genes. Similar to *ARF3/4* in other species, both *MlARF3* and *MlARF4* have two binding sites with sequences complementary to tasiARF (Figure 4A and B). qRT-PCR measurements in 5-mm floral buds showed that in the single and double mutants, *MlARF3* was up-regulated by 1.7~2.5-fold and *MlARF4* was up-regulated by 2.7~3.7-fold (Figure 4C). This moderate up-regulation of *ARF3/ARF4* in the *ago7* and *sgs3* mutant backgrounds is very similar to previous reports in Arabidopsis (Garcia et al., 2006; Hunter et al., 2006), supporting the role of tasiARF in fine-tuning *ARF3/4* expression level.

**Figure 4.**
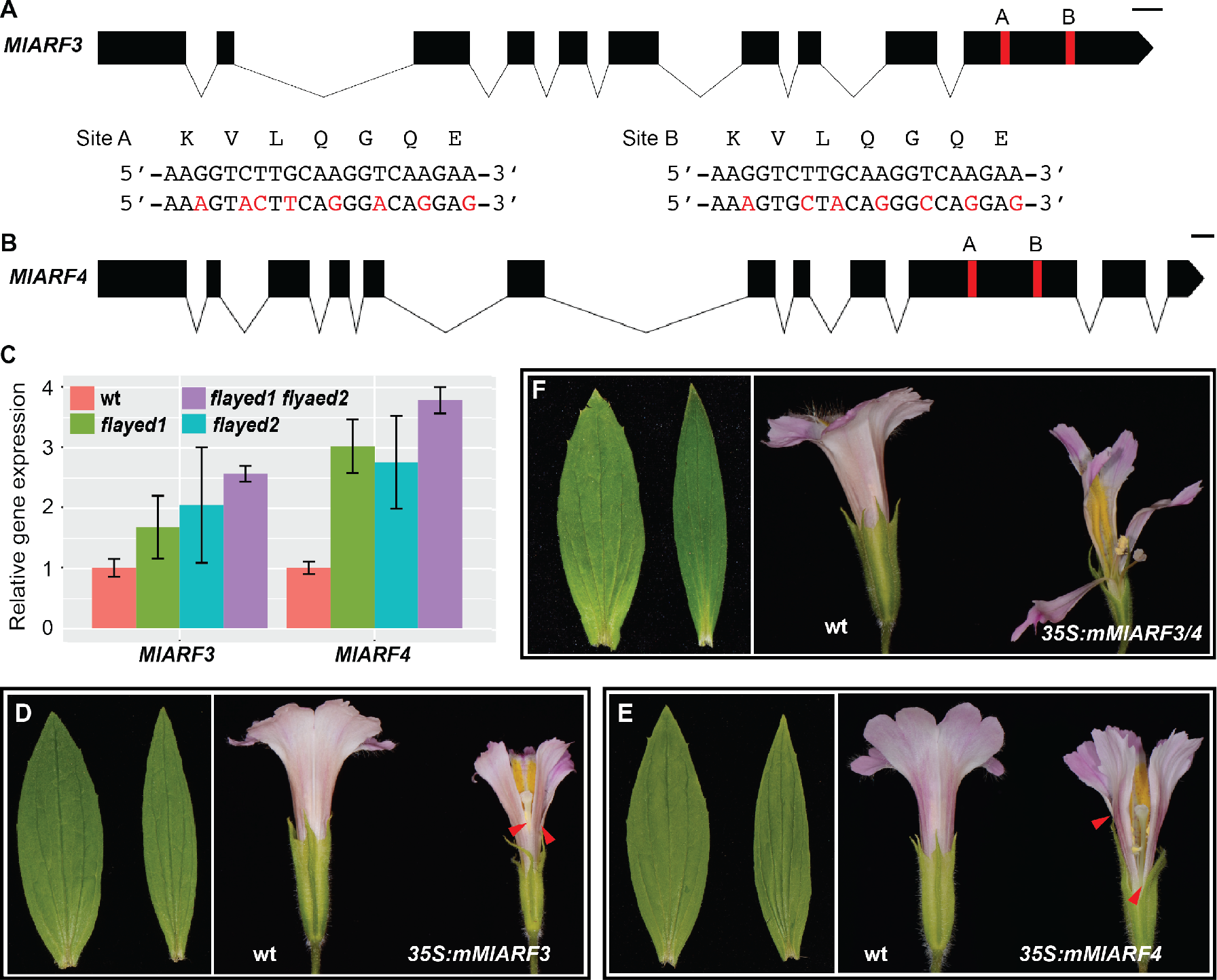
The *flayed1/2* phenotypes are primarily mediated through up-regulation of *MlARF3/4*. (**A** and **B**) Schematics of *MlARF3* (**A**) and *MlARF4* (**B**) gene structure. Red bar: tasiARF binding site. Scale bars are 100 bp. The nucleotides highlighted in red are synonymous substitutions at the two tasiRNA binding sites that were introduced in the *35S:mMlARF3* and *35S:mMlARF4* constructs to circumvent tasiRNA repression. (**C**) Relative transcript level of *MlARF3* and *MlARF4* in 5-mm floral buds as determined by qRT-PCR. *MlUBC* was used as the reference gene. Error bars represent 1 SD from three biological replicates. (**D-F**) Leaf and flower phenotypes of the strongest *35S:mMlARF3* (**D**), *35S:mMlARF4* (**E**), and double transgenic line (**F**). Left: wt; right: transgenic line. The red arrow heads indicate points of petal separation.

To test the third prediction, we transformed the wild-type with a tasiARF-insensitive version of *MlARF3* (*mMlARF3*) and *MlARF4* (*mMlARF4*) with several synonymous substitutions at the tasiARF binding sites (Figure 4A and B), driven by the 35S promoter. We obtained seven independent *35S:mMlARF3* and 14 *35S:mMlARF4* lines. In each case, only two transgenic lines showed obvious phenotypes: their leaves are very similar to the *flayed1/2* mutants (i.e., narrower than the wild-type) and corollas are partially split (indicated by the red arrow heads in Figure 4D and E). qRT-PCR experiments on 5-mm floral buds of the transgenic lines with even the strongest phenotypes showed only moderate overexpression of *MlARF3/4* relative to the wild-type (2~4-fold, Figure S3A and B). Examination of two random *35S:mMlARF4* lines without obvious phenotypes showed no increase in expression level of *MlARF4* (Figure S3C). The lack of *35S:mMlARF3/4* lines with strong transgene expression is in contrast to ectopic expression of *MAGO7* and *MlSGS3* (Figure S1) as well as pigment-related transcription factors in *M. lewisii* (Yuan et al., 2014; Sagawa et al., 2016), where the same 35S promoter could readily drive transgene expression level >10-fold higher than that of the endogenous genes. One possible explanation for this observation is that transgenic lines with very strong *ARF3/4* expression in *M. lewisii* are seedling lethal, as implicated by similar experiments in tomato (Yifhar et al., 2012). Nevertheless, our results show that a moderate up-regulation of *MlARF3/4* can indeed fully recapitulate the leaf phenotype and partially recapitulate the flower phenotype of the *flayed1/2* mutants. Furthermore, a double transgenic line derived from a cross between the strongest *35S:mMlARF3* line and *35S:mMlARF4* line showed dramatic petal fusion defects (Figure 4F). This indicates that *MlARF3* and *MlARF4* may act synergistically in regulating corolla tube formation. Taken together, our results from transgenic manipulation of the *MlARF3/4* expression levels suggest that the *flayed1/2* phenotypes (narrow leaf and split corolla tube) are primarily mediated by the up-regulation of *MlARF3/4*.

### The tasiRNA-ARF Pathway is Required for Preferential Lateral Expansion of the Bases of Petal Primordia and Coordinated Growth of Inter-primordial Regions

To understand how malfunction of the tasiRNA-ARF pathway affects corolla tube formation in *M. lewisii*, we have studied floral organogenesis in the wild-type and the *flayed2* mutant using scanning electron microscopy. Like other species in the lamiid clade (e.g., snapdragon, petunia, morning glory), *M. lewisii* petals are initiated as five separate primordia (Figure 5A). Petal development lags behind stamens in the early stages (Figure 5B and C), but by the time the corolla reaches 0.5 mm in diameter (Figure 5D), petal development progresses rapidly and soon the stamens are found enclosed in the corolla (Figure 5E-H). The developmental stage from 0.3 to 0.4 mm (corolla diameter) is critical for corolla tube formation: during this stage, the bases of the petal primordia quickly expand laterally (to a conspicuously greater extent than the upper portion of the petal primordia; Figure 5M), and the inter-primordial regions also grow coordinately, connecting the initially separate petal primordia. Floral organogenesis of *flayed2* is very similar to that of the wild-type at the early stages (before the corolla reaches 0.3 mm in diameter; Figure 5I). However, during the critical period (0.3~0.4 mm), there is no preferential lateral expansion at the bases of the petal primordia, manifested as the truncate shape of the petal primordium base (Figure 5N), in contrast to the semi-circle shape of the wild-type (Figure 5M). Notably, growth of the inter-primordial regions is also arrested in *flayed2*, leading to a gap between two adjacent petal primordia (Figure 5J-L and indicated by the asterisk in Figure 5N).

**Figure 5.**
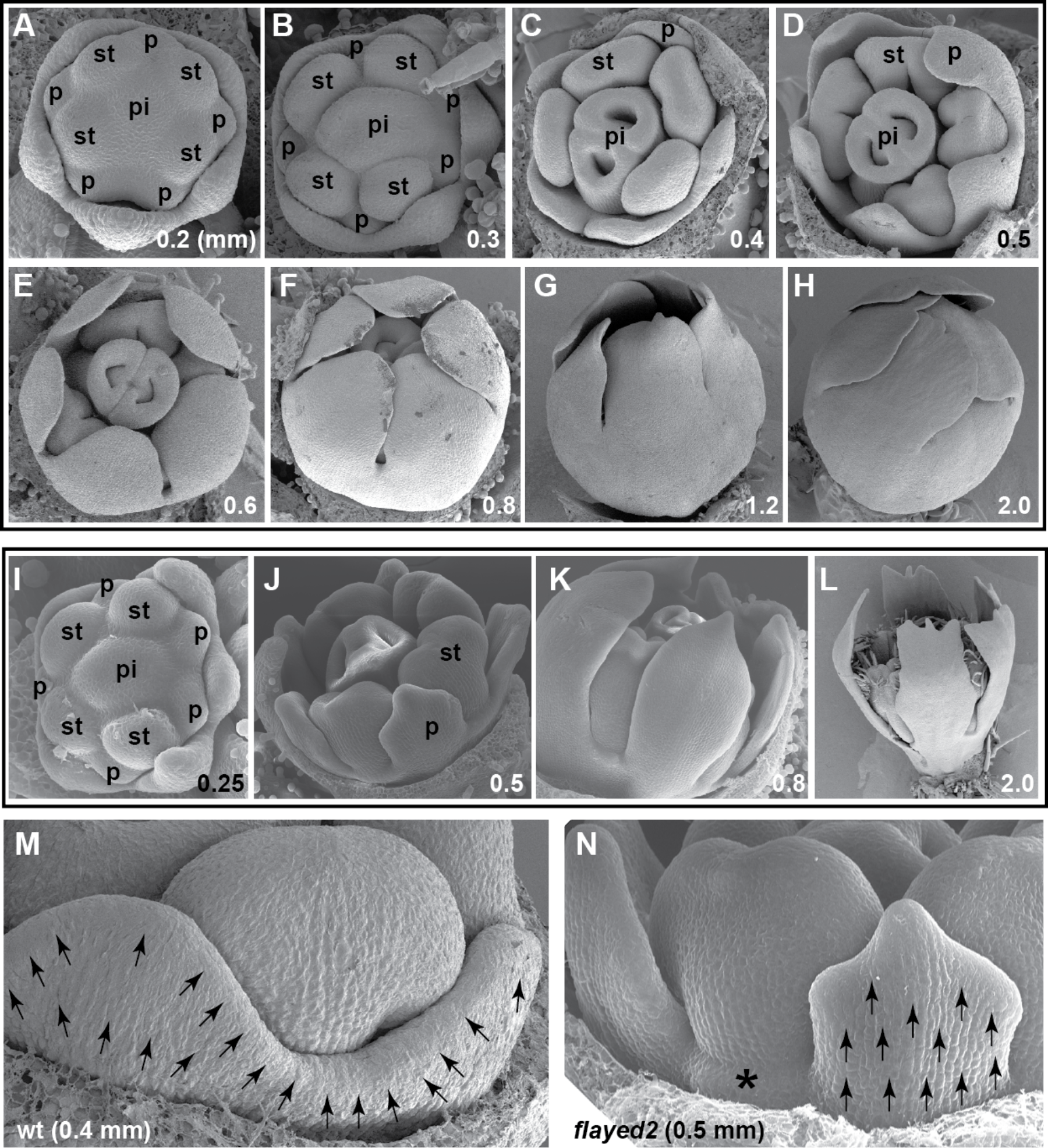
Scanning electron micrographs of *M. lewisii* floral development. (**A-H**) Wild-type LF10. The developmental stages are marked on the bottom right of each image by the diameter (in mm) of the corolla. From 0.4 mm onward, sepals were removed to show the petals. (**I-L**) *flayed2*. (**M** and **N**) Detailed view of two adjacent petal primordia and the inter-primordial region in the wild-type (wt) (**M**) and *flayed2* (**N**). Arrows indicate growth directions, and the asterisk in (N) marks the arrested inter-primordial region. p = petal; st = stamen; pi = pistil.

Given that disruption of tasiRNA biogenesis and the consequent up-regulation of ARF3/4 have been shown to cause reduced lamina growth of lateral organs in multiple plant species (Peragine et al., 2004; Douglas et al., 2010; Yan et al., 2010; Yifhar et al., 2012; Zhou et al., 2013), it is not surprising to observe reduced lateral expansion at the bases of the petal primordia in *flayed2* compared to the wild-type (Figure 5M and N). But how does this relate to the arrest of *upward* growth of the inter-primordial regions?

In a series of careful anatomical studies of various taxa in the Asterids, Nishino recognized that the “co-operation” between the marginal meristem activities of the base of the petal primordia and the upward growth of the inter-primordial regions plays a pivotal role in corolla tube formation (Nishino, 1976, 1978, 1983a, 1983b), although the nature of this “cooperation” was unclear. Considering this earlier insight together with our own results (Figure 5M and N), we speculated that in the wild-type, there is a molecular signal with continuous distribution in the marginal meristematic cells along the petal primordium base and the inter-primordial region, stimulating synchronized growth between the two regions. In the *flayed2* mutant, this growth signal is perhaps reduced or disrupted in spatial distribution, leading to arrested growth in the zone encompassing the margins of primordium base and the inter-primordial region. An obvious candidate for this putative signal is the phytohormone auxin, which is known to promote localized tissue outgrowth and meanwhile suppress organ boundary genes such as *CUP-SHAPED COTYLEDON 1* (*CUC1*) and *CUC2* in Arabidopsis (Bilsborough et al., 2011).

### Auxin Distribution in Developing Corolla Buds Corresponds to Petal Growth Patterns in the Wild-type and Auxin Homeostasis is Altered in *flayed2*

To examine auxin distribution in developing corolla buds, we introduced an auxin reporter construct *DR5rev:mRFPer* (Gallavotti et al., 2008) into the wild-type LF10. In the very early stage where petal primordia just initiate (corresponding to a stage between Figure 5A and B), auxin is accumulated in both petal primordia and inter-primordial regions, with a clear gap between the two (Figure 6A). As the corolla bud reaches 0.4 mm in diameter, auxin becomes concentrated on the apex of the petal primordium and the “synchronized growth zone” encompassing the margins along the primordium base and the inter-primordial region (between the arrow heads in Figure 6B), with relatively weak signal along the upper part of the petal primordium margin. It is worth noting that the auxin localization pattern at this stage corresponds almost perfectly to the petal growth pattern that leads to corolla tube formation (i.e., preferential lateral expansion of the petal primordium base and the coordinated upward growth of the inter-primordial region; Figure 5M). As the corolla bud reaches 0.5 mm in diameter, auxin distribution remains continuous at the base of the petal primordium and the inter-primordial region (demarcated by the arrow heads in Figure 6C), but the gap devoid of auxin between the petal apex and the synchronized growth zone becomes more conspicuous than the 0.4-mm stage. When the corolla bud reaches 0.6 mm in diameter, auxin signal starts to spread more evenly along the entire petal margin (Figure 6D), likely corresponding to later growth of the entire corolla. While all confocal images shown in Figure 6 was taken using *DR5rev:mRFPer* line 2, examination of several additional independent transgenic lines showed very similar results.

**Figure 6.**
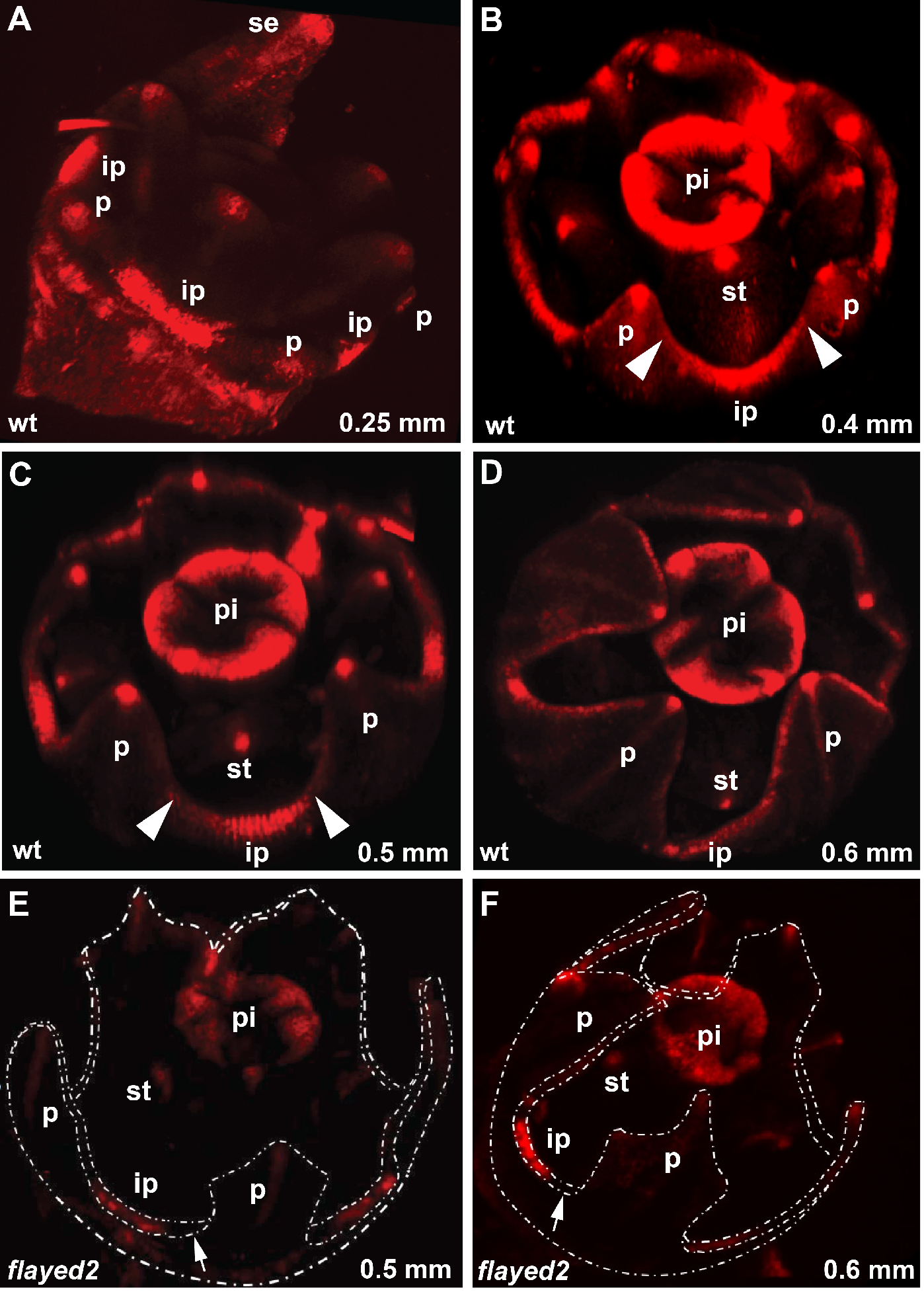
Patterns of auxin distribution in the developing corolla buds of the wild-type and *flayed2*, as reflected by the *DR5rev:mRFPer* reporter signal. (**A-D**) Wild-type (wt). The developmental stages are marked on the bottom right of each image by the diameter (in mm) of the corolla. The white arrow heads in (**B**) and (**C**) demarcate the synchronized growth zone encompassing the marginal meristematic cells at the base of the petal primordia and the inter-primordial cells. se = sepal; p = petal; ip = inter-primordial region; st = stamen; pi = pistil. (**E** and **F**) *flayed2*. Shown in the same style as the wild-type. The overall fluorescence signal is much weaker in the mutant compared to the wild-type. Petal primordia and inter-primordial regions are outlined in white dashed lines. White arrows mark the gap at the junction between the petal primordium base and the inter-primordial region.

To test whether auxin homeostasis is altered in the *flayed2* mutant, we crossed *DR5rev:mRFPer* line 2 with *flayed2*, and analyzed F_2_ individuals that are homozygous for the *flayed2* mutation and with the *DR5rev:mRFPer* transgene. When imaged under the same conditions, the *flayed2* corolla buds showed much weaker auxin signal overall than the wild-type (Figure 6E and F). In particular, the junction between the petal primordium base and the inter-primordial region (arrows in Figure 6E and F) showed no signal at all, consistent with the lack of synchronized growth between these two regions in the mutant (Figure 5N). The decreased auxin signals in *flayed2* is likely caused by up-regulation of *MlARF3/4*, as ARF3 has been implicated in auxin homeostasis in Arabidopsis by directly repressing auxin biosynthesis and transport (Simonini et al., 2017).

One potential consequence of the reduced and discontinuous auxin distribution in the “synchronized growth zone” of *flayed2* corolla buds is up-regulation of organ boundary genes such as *NAM/CUC1/CUC2* homologs (Bilsborough et al., 2011). To test this possibility, we mined an unpublished transcriptome assembled from various developmental stages of *M. lewisii* floral buds, to search for genes that are closely related to *NAM/CUC1/CUC2*, which can be identified using the amino acid motif “HVSCFS[N/T/S]” downstream of the NAC domain (Ooka et al., 2003) in addition to overall sequence identity. The “HVSCFS[N/T/S]” motif also corresponds to the recognition site for miR164 (Laufs et al., 2004; Mallory et al., 2004). We found three genes named *MlNAC1/2/3* with highest sequence identity to *NAM/CUC1/CUC2* and with the putative miR164 recognition site (Figure S4A and B). qRT-PCR experiments on 2-mm floral buds showed no expression difference between the wild-type and *flayed2* for *MlNAC1* or *MlNAC2;* however, *MlNAC3* transcript level is ~8-fold higher in *flayed2* than in the wild-type (Figure S4C). The functional relevance of *MlNAC3* up-regulation in corolla tube formation will need be tested by future transgenic experiments.

## DISCUSSION

This study represents the first step of a systematic effort towards understanding the developmental genetics of corolla tube formation using the new model system *Mimulus lewisii*. We show that the tasiRNA-ARF pathway is required for the synchronized growth between the petal primordium base and the inter-primordial region at early stages of corolla development. Furthermore, we discovered that auxin distribution is continuous in this synchronized growth zone in the wild-type, but becomes much reduced in the *flayed2* mutant with split corolla tubes, explaining the inhibition of lateral expansion at the base of the petal primordia and complete arrest of the upward growth of the inter-primordial regions observed in the mutant. Together, these results suggest a new conceptual model for the developmental genetic control of corolla tube formation.

### Role of the tasiRNA-ARF Pathway in Petal Fusion

Although this is the first detailed study investigating the role of the tasiRNA-ARF pathway in corolla tube formation, a previous study on leaf development in the family Solanaceae has mentioned in passing that malfunction of the tasiRNA-ARF pathway led to unfused corolla in tomato and tobacco flowers (Yifhar et al., 2012). Like *Mimulus*, the family Solanaceae belongs to the asterid clade. This suggests that the role of the tasiRNA-ARF pathway in corolla tube formation is likely conserved across asterid plants. More surprisingly, two other studies on leaf development in *Medicago truncatula* and *Lotus japonicus*, both belonging to the legume family (Fabaceae), also implicated an indispensable role of the tasiRNA-ARF pathway in petal fusion (Yan et al., 2010; Zhou et al., 2013). Typical legume flowers have separate dorsal and lateral petals; but the two ventral petals often fuse into a prow-shaped structure (i.e., the “keel”). In the *ago7* mutants of *Medicago truncatula* and *Lotus japonicus*, the two ventral petals become separated. The family Fabaceae belongs to the clade Rosids. Given that the vast majority of rosid species (e.g., *Arabidopsis thaliana*) produce flowers with completely separate petals (i.e., polypetalous), with only a few exceptions in derived lineages (Zhong and Preston, 2015), it is clear that the tasiRNA-ARF pathway was independently recruited to enable petal fusion in the legume species.

There are two significant insights emerging from our molecular and phenotypic analyses of the tasiRNA-ARF pathway in *Mimulus* that were not known from the aforementioned studies that focused on leaf development: (i) The tasiRNA-ARF pathway is required not only for the lamina growth of the petal primordia, but also for the upward growth of the inter-primordial regions (Figure 5M and N). In fact, we think that the synchronized growth between the petal primordium base and the inter-primordial region is the key to corolla tube formation; (ii) There is a tight correlation between auxin distribution and the growth pattern in developing corolla buds (Figure 5M; Figure 6B and C). The drastic reduction of auxin concentration in the *flayed2* mutant compared to the wild-type is likely due to up-regulation of *MARF3/4*, as there are evidence from Arabidopsis that ARF3 can directly repress auxin biosynthesis and transportation (Simonini et al., 2017).

Given the well-known function of the tasiRNA-ARF pathway in lamina growth of lateral organs (Fahlgren et al., 2006; Garcia et al., 2006; Hunter et al., 2006; Nagasaki et al., 2007; Nogueira et al., 2007; Douglas et al., 2010 Yan et al., 2010; Yifhar et al., 2012; Zhou et al., 2013) and the reduced petal width of the *flayed1/2* mutants, it is easy to over-interpret the significance of lamina growth of the petal primordia in the “fusion” of adjacent petals. However, the following observations suggest that lamina growth of the petal primordia *per se* is not the key to corolla tube formation: (**i**) In some of the *35S:MlAGO7* and *35S:MlSGS3* transgenic lines (in the corresponding mutant background), petal width was restored to wild-type level, but the petals remained unfused (Figure 2C and D). In addition, we have several other yet-to-be-characterized *flayed* mutants with split corolla but normal petal width. These observations suggest that petal lamina growth at the wild-type level is not sufficient for tube formation; (**ii**) Petal width of a previously characterized *M. lewisii* mutant, *act1-D*, is very similar to that of *flayed1/2* (Ding et al., 2017), but the corolla tube of *act1-D* is intact. This suggests that reduced petal lamina growth does not necessarily affect petal fusion. Instead, we think it is the synchronization between the lateral expansion of the petal primordium base and the upward growth of the inter-primordia region, most likely directed by auxin signaling, that holds the key to corolla tube formation.

The difference in auxin accumulation patterns between *flayed2* and the wild-type (Figure 6) suggests that the role of the tasiRNA-ARF pathway in corolla tube formation lies in the regulation of auxin homeostasis. But how exactly does the tasiRNA-ARF pathway regulates the spatial pattern and dynamics of auxin distribution in the developing corolla buds of *Mimulus* remains to be elucidated.

### A New Conceptual Model for the Developmental Genetic Control of Corolla Tube Formation

Two recent attempts of building a conceptual framework for floral organ fusion in general (Specht and Howarth, 2015) or petal fusion in particular (Zhong and Preston, 2015) both emphasized the genetic regulatory network underlying organ boundary formation and maintenance. The rationale for such emphasis was explicitly stated by Specht and Howarth (Specht and Howarth, 2015): “*fusion as a process may more accurately be defined as a lack of organ separation*”. While these attempts represent an important step towards a mechanistic understanding of the developmental process of corolla tube formation, to some degree they have neglected the insight provided by earlier morphological and anatomical studies (i.e., the importance of the “co-operation” between the rapid lateral expansion of the petal primordium base and the upward growth of the inter-primordial region), and have not provided a logical explanation to the previously reported corolla tube mutants (e.g., morning glory *feathered*, petunia *maewest*) (Iwasaki and Nitasaka, 2006; Vandenbussche et al., 2009) in terms of organ boundary formation or maintenance. Overemphasis on the “lack of organ separation” may represent an underestimation of the complexity of the corolla tube formation process, with the implication that simple loss-of-function mutations in some organ boundary genes in polypetalous species (e.g., *Arabidopsis thaliana*) could produce a functional corolla tube. As far as we are aware of, despite extensive mutagenesis of Arabidopsis in the past 40 years, such a mutant has not been reported.

Integrating our results on the tasiRNA-ARF pathway and auxin distributions in *Mimulus lewisii* with previous reports of corolla tube mutants (Iwasaki and Nitasaka, 2006; Vandenbussche et al., 2009) as well as historical insights from anatomical studies (Nishino, 1978, 1983a, 1983b), we propose a new conceptual model for the developmental genetic control of corolla tube formation (Figure 7). At the heart of the model is auxin-induced synchronized growth of the marginal meristematic cells at the base of the petal primordia and the inter-primordial cells, providing a molecular explanation for the “co-operation” between the petal primordium base and the inter-primordial region observed in anatomical studies. Upstream of this core module is the genetic regulatory network controlling adaxial/abaxial polarity and lamina growth of lateral organs (Nakata and Okada, 2013; Tsukaya, 2013; Kuhlemeier and Timmermans, 2016), which is conserved in a wide range of angiosperms and is somehow coopted in sympetalous species to regulate auxin homeostasis in the synchronized growth zone. Downstream of this core module lie the organ boundary genes that would suppress localized tissue growth if not repressed by auxin signaling.

**Figure 7.**
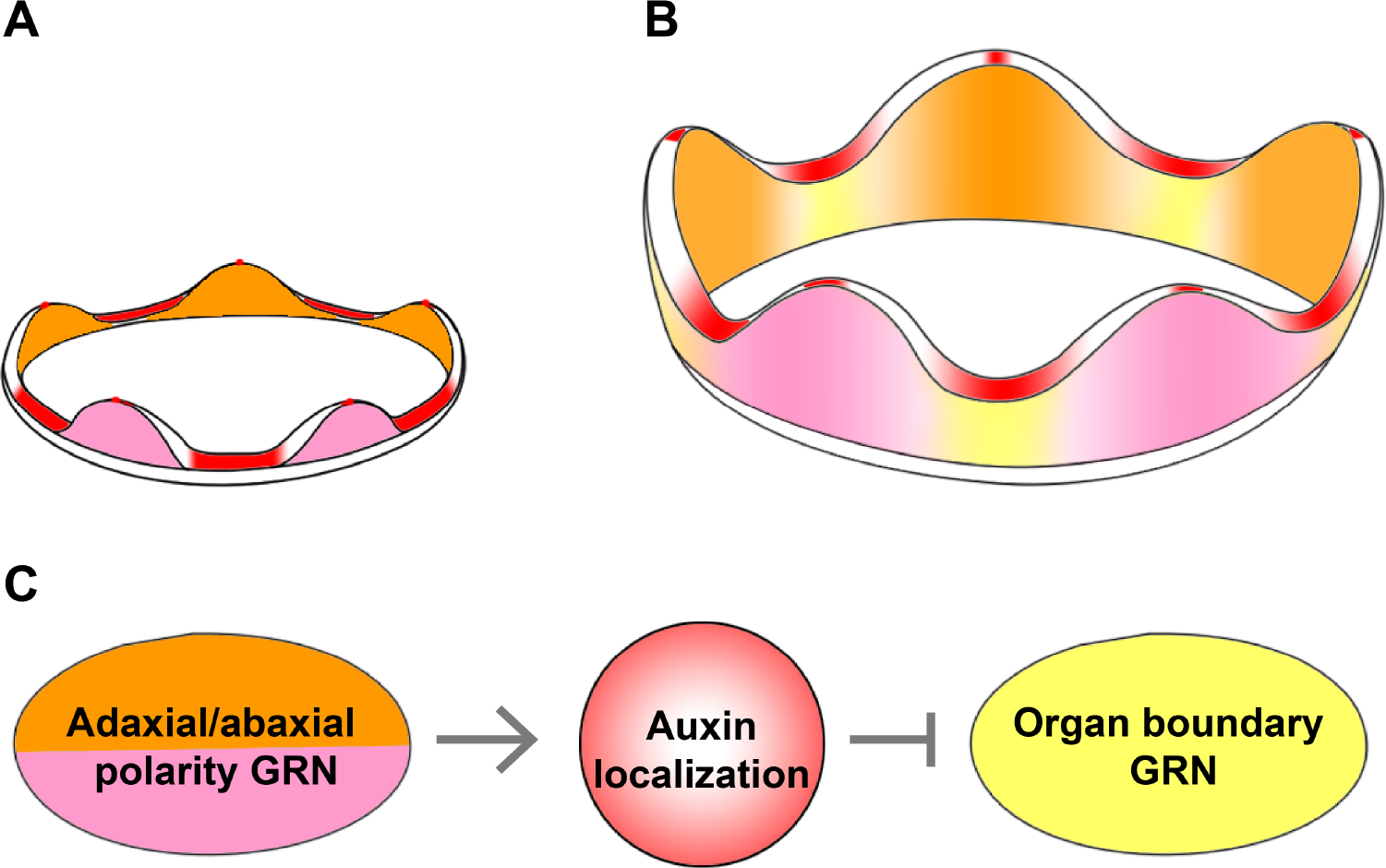
A conceptual model for the developmental genetic control of corolla tube formation in *Mimulus lewisii*. (**A** and **B**) Schematics of auxin distribution patterns at very early stage when petal primordia just initiate (**A**) and when the corolla tube is formed by synchronized growth of the petal primordium base and the inter-primordial region (**B**). Pink and orange color indicates the abaxial and adaxial side of the petal, respectively. Yellow color indicates the inter-primordial region. (**C**) Genetic relationships among the three hypothetical modules. GRN: Genetic Regulatory Network.

This model can readily explain the phenotypes of loss-of-function mutations in the morning glory *FEATHERED* gene and the petunia *MAEWEST* gene, which encode KANADI and WOX transcription factors, respectively (Iwasaki and Nitasaka, 2006; Vandenbussche et al., 2009). Together with the tasiRNA-ARF pathway, these transcription factors are part of the genetic network regulating leaf adaxial-abaxial polarity and lamina growth (Nakata and Okada, 2013; Tsukaya, 2013; Kuhlemeier and Timmermans, 2016). Recent studies in other model systems such as *Arabidopsis, Medicago*, and *Nicotiana*, have demonstrated that these polarity regulators largely function by modulating auxin homeostasis (Tadege et al., 2011; Huang et al., 2014; Simonini et al., 2017). According to our model, interfering with this genetic regulatory network is expected to reduce or disrupt the continuous auxin distribution in the synchronized growth zone, as shown in the *flayed2* mutant (Figure 5N; Figure 6E and F), and consequently resulting in unfused corolla.

Our model also provides a plausible explanation for the petal fusion defects observed when a chimeric repressor of AtTCP5 was over-expressed in the Japanese morning glory (Ono et al., 2012). Chimeric repressors of CIN-like TCP transcription factors, including TCP5 in Arabidopsis, are known to activate organ boundary genes such as *CUC1/2* (Koyama et al., 2007), which are otherwise suppressed by auxin (Bilsborough et al., 2011; Figure S4C). Ectopic activation of *CUC1/2* is expected to prevent the upward growth of the inter-primordial regions (i.e., boundary between adjacent petal primordia). Also consistent with the “organ boundary” module (Figure 7C) is a recent observation in snapdragon — the expression of the *CUC* ortholog, *CUPULIFORMIS*, is cleared from the inter-primordial regions shortly after petal initiation but later is reactivated in the sinuses between adjacent corolla lobes (Rebocho et al., 2017). Through computational modeling, Rebocho et al. (2017) showed that this “gap” of *CUPULIFORMIS* expression (between the base of the corolla and the sinuses) is necessary for corolla tube formation.

In addition to explaining these previous observations, our model predicts that mis-regulation of auxin biosynthesis, polar transport, and signaling within and between petal primordia, in transgenic or mutant plants of sympetalous species should result in defective corolla tubes. It also predicts that transgenic manipulation of other components of the leaf polarity/lamina growth network (e.g., *AS1, AS2, HD-ZIPIII*) or ectopic expression of organ boundary genes (e.g., *CUC*, *ORGANBOUNDARY1, JAGGED LATERAL ORGAN*) (Aida et al., 1997; Borghi et al., 2007; Cho and Zambryski, 2011) in sympetalous species may also result in unfused corollas. The availability of multiple corolla tube mutants, the ease of bulk segregant analysis to identify mutant genes, and the amenability of *Agrobacterium*-mediated *in planta* transformation make *Mimulus* a favorable system to test these predictions and to dissect the detailed molecular mechanisms and developmental process of corolla tube formation.

## METHODS

### Plant Materials and Growth Conditions

EMS mutagenesis was performed using the *Mimulus lewisii* Pursh inbred line LF10 as described (Owen and Bradshaw, 2011). Another inbred line SL9 was used to generate the *“flayed* × SL9” F_2_ populations. Plants were grown in the University of Connecticut research greenhouses under natural light supplemented with sodium vapor lamps, ensuring a 16-hr day length.

### Phenotypic Characterization

To quantify phenotypic differences between the mutants and wild-type, we measured the widths of the dorsal, ventral, and lateral petals using a digital caliper. We also measured the lengths and widths of the fourth leaf (the largest leaf) of mature plants. To further evaluate whether the width difference is due to change in cell number, cell size or both, width of the abaxial epidermal cells of the dorsal petal lobe was measured following a previously described procedure (Ding et al., 2017).

### Genomic Analyses for Causal Gene Identification

To identify causal genes underlying *flayed1* and *flayed2*, we employed a hybrid strategy that combines the advantages of bulk segregant analysis and genome comparisons between multiple EMS mutants, as described previously (LaFountain et al., 2017). Briefly, for each mutant an F_2_ population was produced by crossing the homozygous mutant (generated in the LF10 background) and the mapping line SL9. DNA samples from 96 F_2_ segregants displaying the mutant phenotype (i.e., homozygous for the causal mutation) were pooled with equal representation. A small-insert library was then prepared for the pooled sample and was sequenced using an Illumina HiSeq 2000 platform at the University of North Carolina High Throughput Sequencing Facility. About 213 and 448 million 100-bp paired-end reads were generated for *flayed1* and *flayed2*, respectively.

The short reads were mapped to the LF10 genome assembly version 1.8 (http://monkeyflower.uconn.edu/resources/) using CLC Genomics Workbench 7.0 (Qiagen, Valencia, CA). The causal mutation should be: (**i**) homozygous for the pooled sample (i.e., 100% SNP frequency in the “F_2_ reads - LF10 genome” alignment); and (**ii**) unique to each mutant (i.e., any shared 100% SNPs between mutants are most likely due to assembly error in the reference genome or non-specific mapping of repetitive sequences). After comparisons to the SNP profiles of previously published mutants, *guideless* (Yuan et al., 2013b), *rcp1* (Sagawa et al., 2016), *act1-D* (Ding et al., 2017), and *rcp2* (Stanley et al., 2017), we narrowed the causal mutation to 39 and 19 candidate SNPs for *flayed1* and *flayed2*, respectively (Figure S5; Table S2 and S3).

### Small RNA Sequencing and Analyses

For small RNA sequencing, total RNA was first extracted using the Spectrum Plant Total RNA Kit (Sigma-Aldrich) from 5-mm floral buds of LF10, *flayed1*, and *flayed2* (two biological replicates for each genotype). Small RNA libraries were then constructed using the Illumina TruSeq Small RNA Sample Preparation Kits, with the total RNA as starting material. The libraries were sequenced on an Illumina HiSeq 2500 at the Delaware Biotechnology Institute (Newark, DE). Small RNA reads were quality-controlled and adaptor-trimmed before calculating tasiRNA abundance, as described in Xia et al. (2017).

### Quantitative RT-PCR

RNA extraction and cDNA synthesis were as previously described (Yuan et al., 2013a). cDNA samples were diluted 10-fold before qRT-PCR. All qRT-PCRs were performed using iQ SYBR Green Supermix (Bio-Rad) in a CFX96 Touch Real-Time PCR Detection System (Bio-Rad). Samples were amplified for 40 cycles of 95 °C for 15 s and 60 °C for 30 s. Amplification efficiencies for each primer pair were determined using critical threshold values obtained from a dilution series (1:4, 1:8, 1:16, 1:32) of pooled cDNA. *MlUBC* was used as a reference gene as described (Yuan et al., 2013a). Primers used for qRT-PCR are listed in Table S4.

### Plasmid Construction and Plant Transformation

To generate the *35S:MlAGO7* and *35S:MlSGS3* constructs for the rescue experiments, we first amplified the full-length CDS of *MlAGO7* and *MlSGS3* from the wild-type LF10 cDNA using the Phusion enzyme (NEB, Ipswich, MA). For each gene, the amplified fragment was cloned into the pENTR/D-TOPO vector (Invitrogen) and then a linear fragment containing the CDS flanked by the attL1 and attL2 sites was amplified using M13 primers. This linear fragment was subsequently recombined into the Gateway vector pEarleyGate 100 (Earley et al., 2006), which drives transgene expression by the CaMV *35S* promoter. To generate the *35S:mMlARF3/4* constructs, CDS of sRNA insensitive forms of *MARF3* (*mARF3*) and *MlARF4* (*mARF4*) that carries synonymous substitutions in the two tasiRNA recognition sites were synthesized by GenScript (NJ, USA) and then cloned into the pEarleyGate 100 destination vector as described for the *35S:MlAGO7* and *35S:MlSGS3* constructs. All plasmids were verified by sequencing before being transformed into *Agrobacterum tumefaciens* strain GV3101 for subsequent plant transformation, as described in Yuan et al. (2013a). Primers used for plasmid construction and sequencing are listed in Table S5.

#### Scanning Electron Microscopy

Flower buds were fixed overnight in Formalin-Acetic-Alcohol (FAA) at 4°C, dehydrated for 30 min through a 50%, 60%, 70%, 95%, and 100% alcohol series. Samples were then critical-point dried, mounted, and sputter coated before being observed using a NOVA NanoSEM with Oxford EDX at 35 kV at UConn’s Bioscience Electron Microscopy Laboratory.

### Confocal Microscopy

Early developing floral buds (with sepals removed if necessary) carrying the *DR5rev:mRFPer* construct were laser scanned in red channel with a *Z*-stack. All fluorescence images were acquired using a Nikon A1R confocal laser scanning microscope equipped with a 20X objective (Plan APO VC 20X/1.20) at UConn’s Advanced Light Microscopy Facility.

## Accession Numbers

Short read data have been deposited in NCBI SRA (BioProject PRJNA423263); small RNA data have been deposited in NCBI GEO (GSE108530); annotated gene sequences have been deposited in GenBank (MG669632-MG669634 and MF084285).

## ACKNOWLEDGEMENTS

We are grateful to Dr. Toby Bradshaw (University of Washington) for encouragement and initial support for generating the bulk segregant data in his laboratory. We thank Clinton Morse, Matt Opel, and Adam Histen for plant care in the UConn EEB Research Greenhouses. The *DR5rev:mRFPer* plasmid was kindly provided by Dr. David Jackson. This work was supported by the University of Connecticut start-up funds and an NSF grant (IOS-1558083) to Y-W.Y., and an NSF grant (IOS-1257869) to B.C.M.

## AUTHOR CONTRIBUTIONS

Y.W.Y. conceived the project. B.D. and Y.W.Y. designed the research. B.D., R.X., Q.L., V.G., J.M.S., L.E.S., M.S., and P.K.D. performed the experiments. B.D., Y.W.Y., P.K.D., R.X., and B.C.M. analyzed data. B.D. and Y.W.Y. wrote the manuscript with input from all the authors.

## Supporting Information

**Figure S1.**
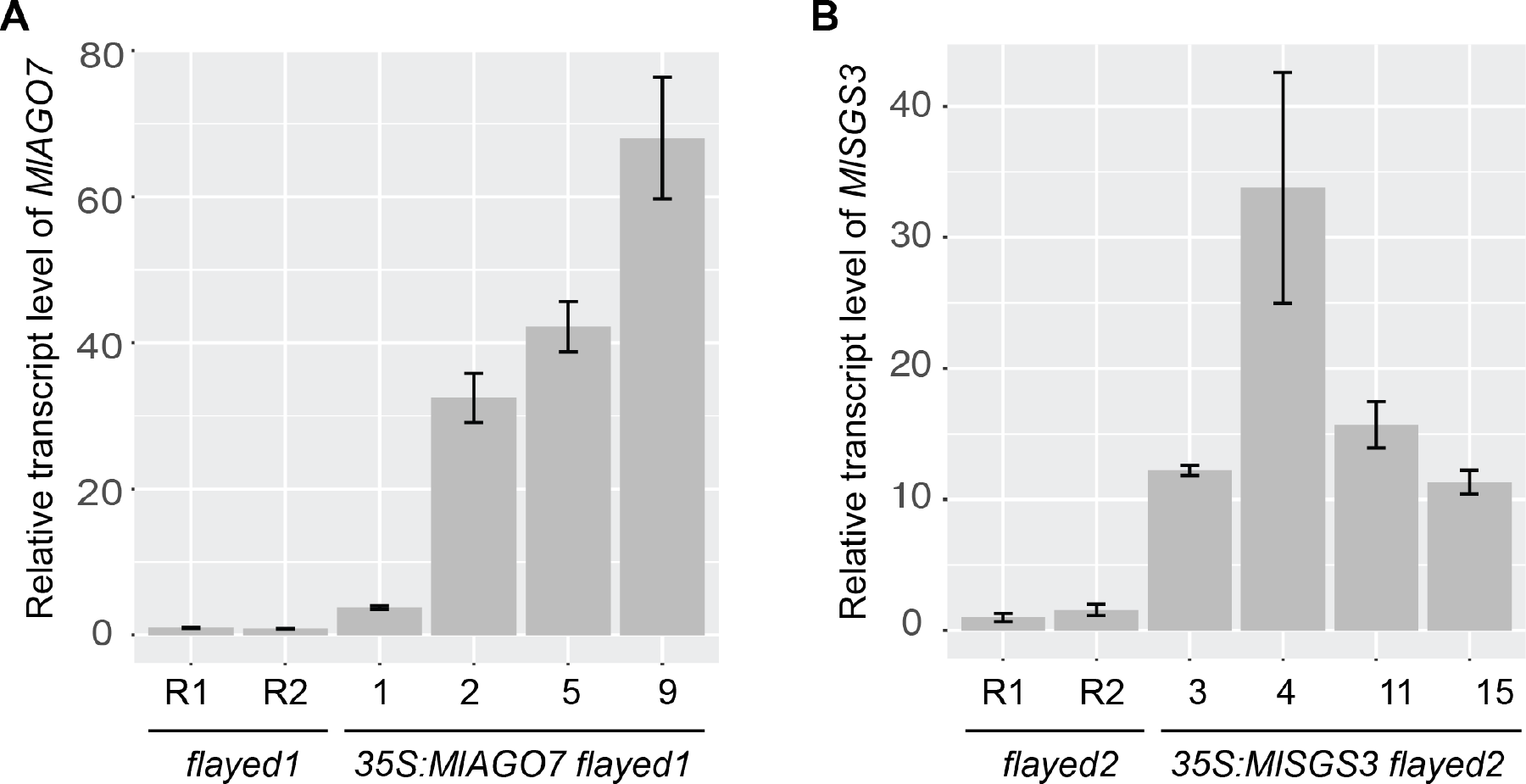
Transgene expression level in the rescue lines. Relative transcript level of *MIAGO7* **(A)** and *MISGS3* **(B)** in 5-mm floral buds of four representative, fully rescued over-expression lines compared to the corresponding mutant backgrounds, as determined by qRT-PCR. R1 and R2 represent two biological replicates. *MlUBC* was used as the reference gene. Error bars represent 1 SD from three technical replicates.

**Figure S2.**
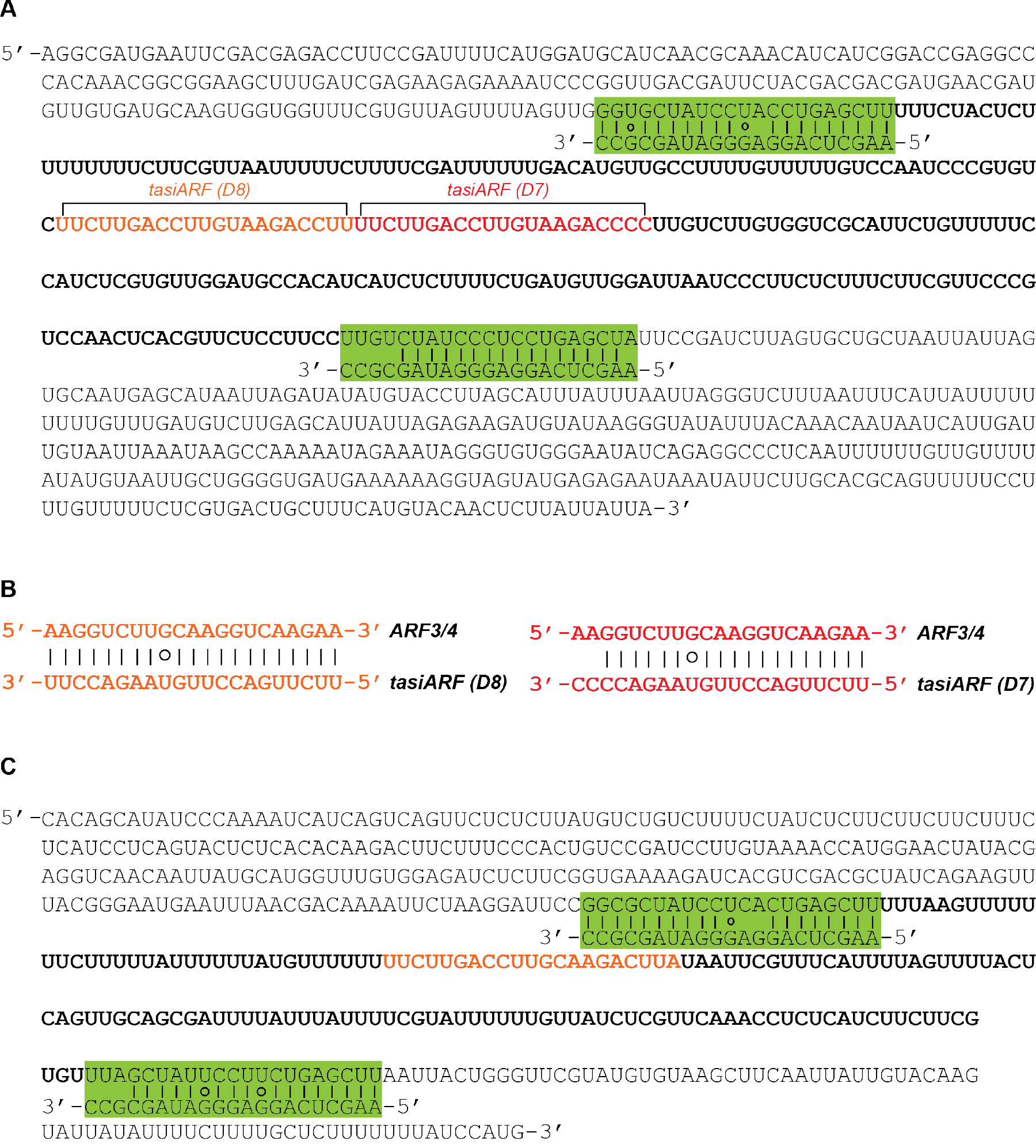
Annotation of the *TAS3* transcripts in *Mimulus lewisii*. **(A)** *TAS3L*. The two miR390 binding sites are indicated by the green boxes; the tandem tasiARFs are highlighted in orange and red fonts. **(B)** Sequence complementarity between the two tasiARFs and the tasiARF-binding sites of *MlARF3/4*. **(C)** *TAS3S*. Highlighted are the miR390 binding sites and the single tasiARF sequence.

**Figure S3.**
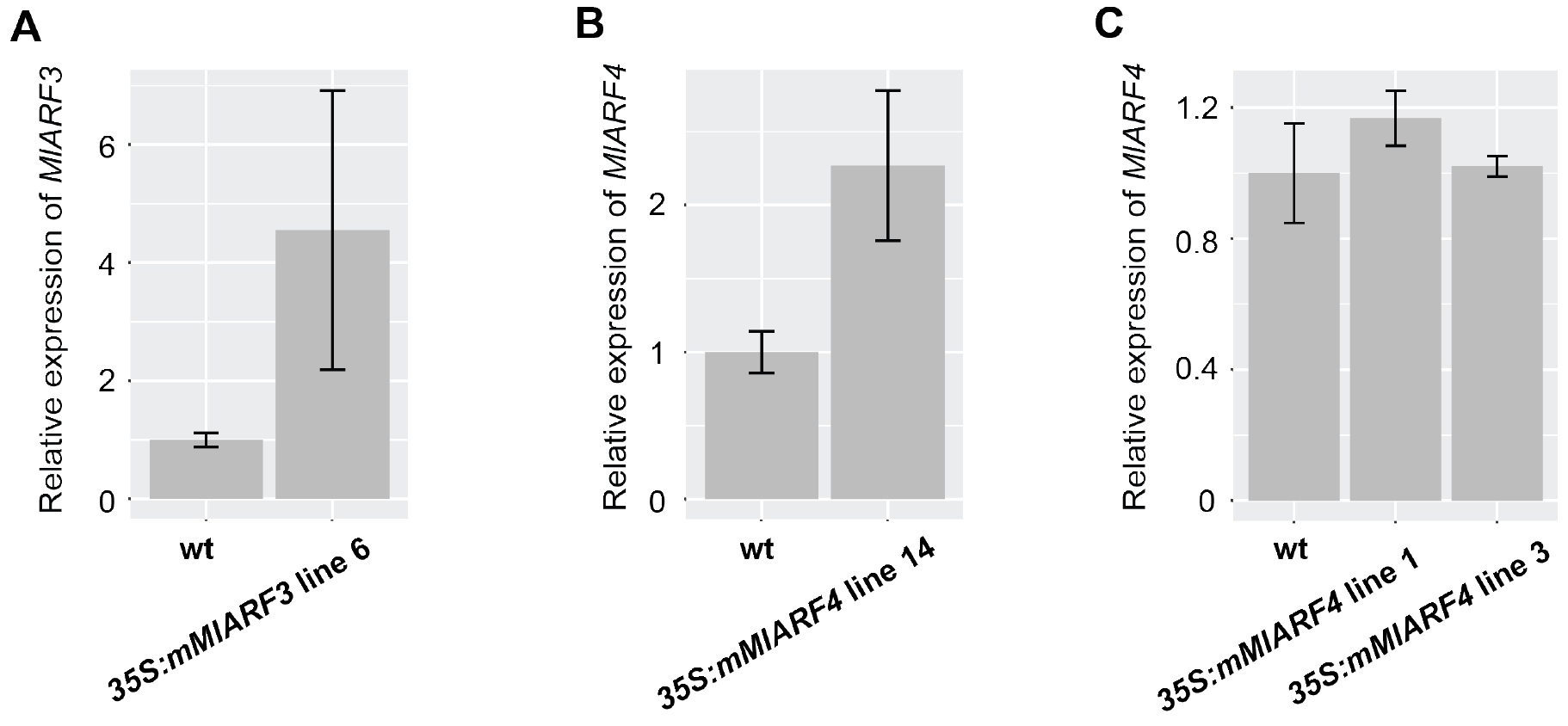
Relative transcript level of *MIARF3/4* in 5-mm floral buds as determined by qRT-PCR. **(A)** *MIARF3* in the *35S:mMlARF3* line with the strongest phenotype. **(B)** *MIARF4* in the *35S:mMlARF4* line with the strongest phenotype. **(C)** *MIARF4* in two *35S:mMlARF4* transgenic lines without any obvious phenotypes. *MlUBC* was used as the reference gene. Error bars represent 1 SD from three biological replicates.

**Figure S4.**
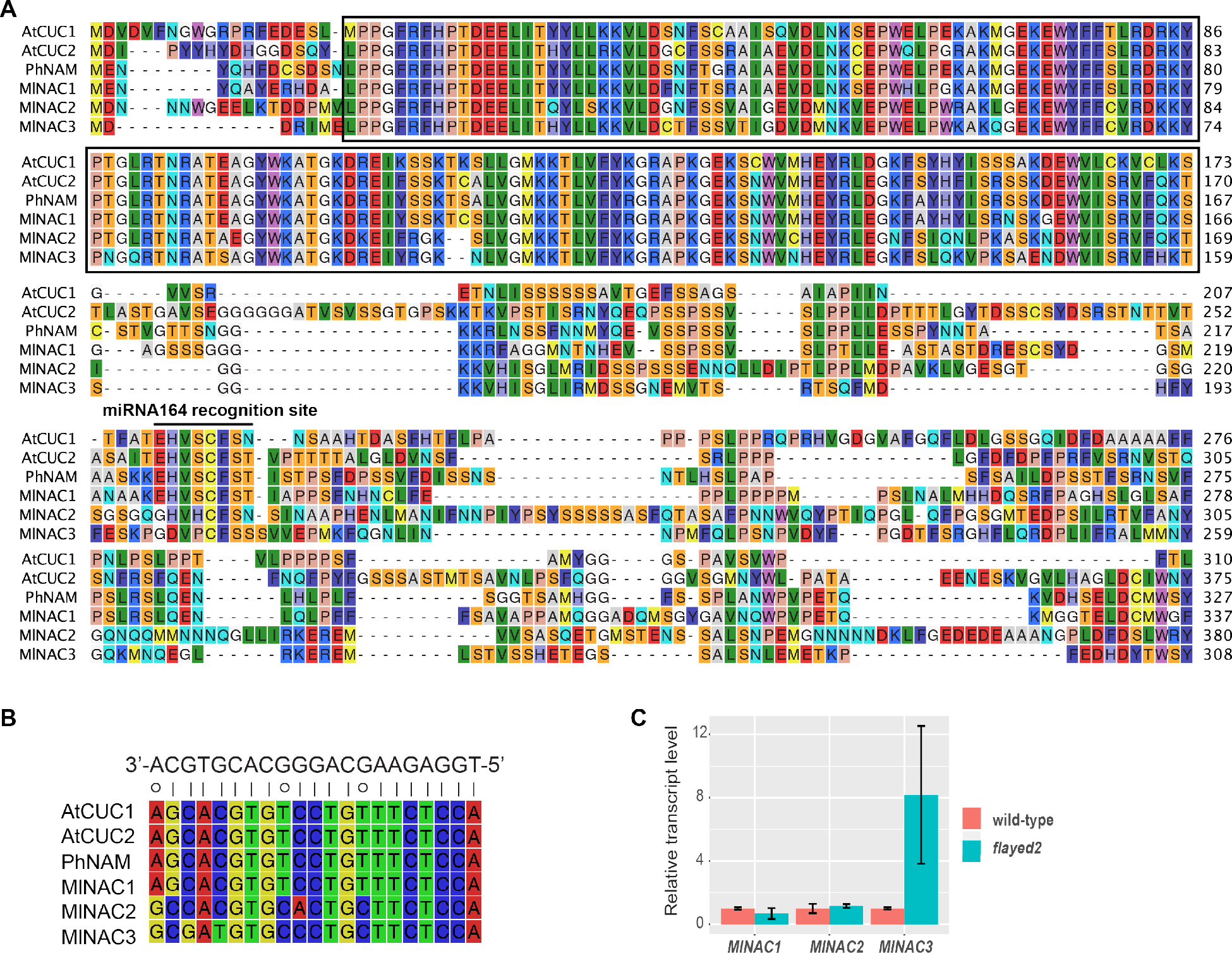
A *NAM/CUC1/2* related gene is up-regulated in *flayed2* compared to the wild-type. **(A)** Amino acid alignment of Arabidopsis CU1/2 and petunia NAM with three *M. lewisii* homologs. Box indicates the NAC domain and the horizontal bar indicates the amino acid motif “HVSCFS[N/S/T]”. **(B)** Nucleotide sequence alignment of the miR164 recognition site. **(C)** Relative transcript level of *MlNAC1/2/3* in 2-mm floral buds as determined by qRT-PCR. *MlUBC* was used as the reference gene. Error bars represent 1 SD from four biological replicates.

**Figure S5.**
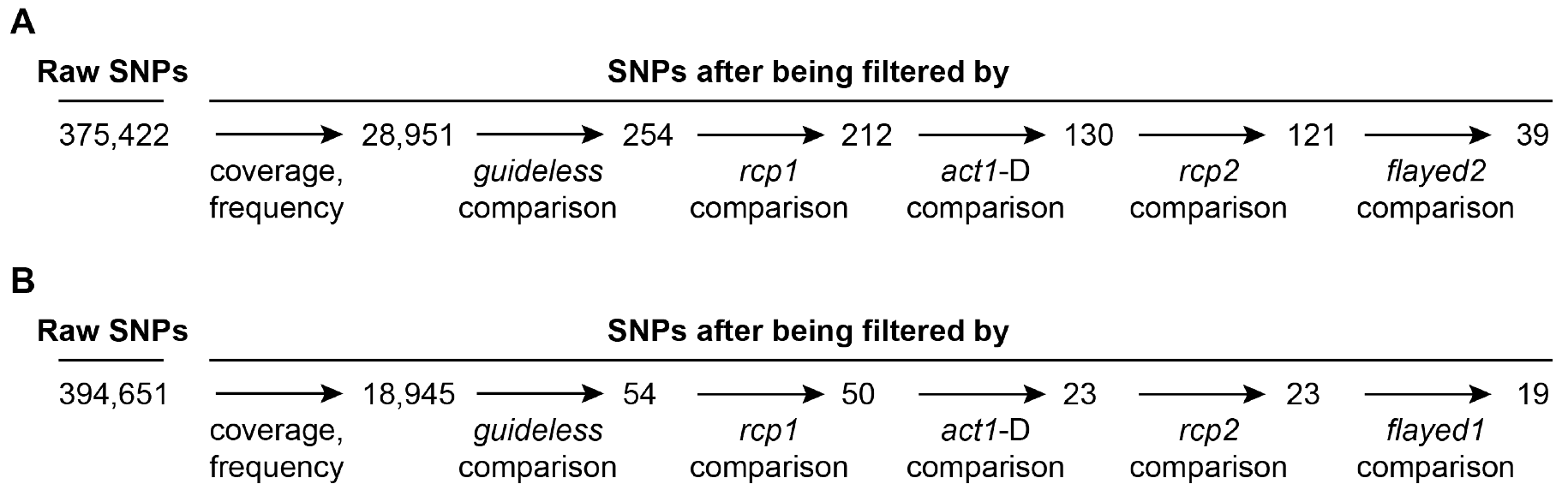
SNP profile comparison between different EMS mutants. **(A)** Comparison between *flayed1* and other mutants. **(B)** comparison between *flayed2* and other mutants.

**Table S1.** Measurements of width (mm) of the dorsal, lateral and ventral petal, and the length (mm) and width (mm) of the fourth leaf in *Mimulus lewisii* wild-type LF10 (n = 18), *flayed1* (n = 10), *flayed2* (n = 12) and the double mutant (n = 12) (mean±SD).

| Trait | Wild-type | *flayed1* | *flayed2* | *flayed1 flayed2* |
| --- | --- | --- | --- | --- |
| Dorsal Petal Width | 12.71±1.13 | 8.88±0.63 | 8.23±0.32 | 8.13±0.37 |
| Lateral Petal Width | 9.7±0.73 | 6.92±0.51 | 6.80±0.39 | 6.44±0.49 |
| Ventral Petal Width | 9.68±0.84 | 6.1±0.45 | 6.29±0.32 | 6.02±0.28 |
| Leaf Width | 34.78±2.02 | 24±1.7 | 23±1.53 | 24.33±1.56 |
| Leaf Length | 92.31±4.02 | 91.6±6.74 | 93.33±4.70 | 97.75±7.5 |

**Table S2.**
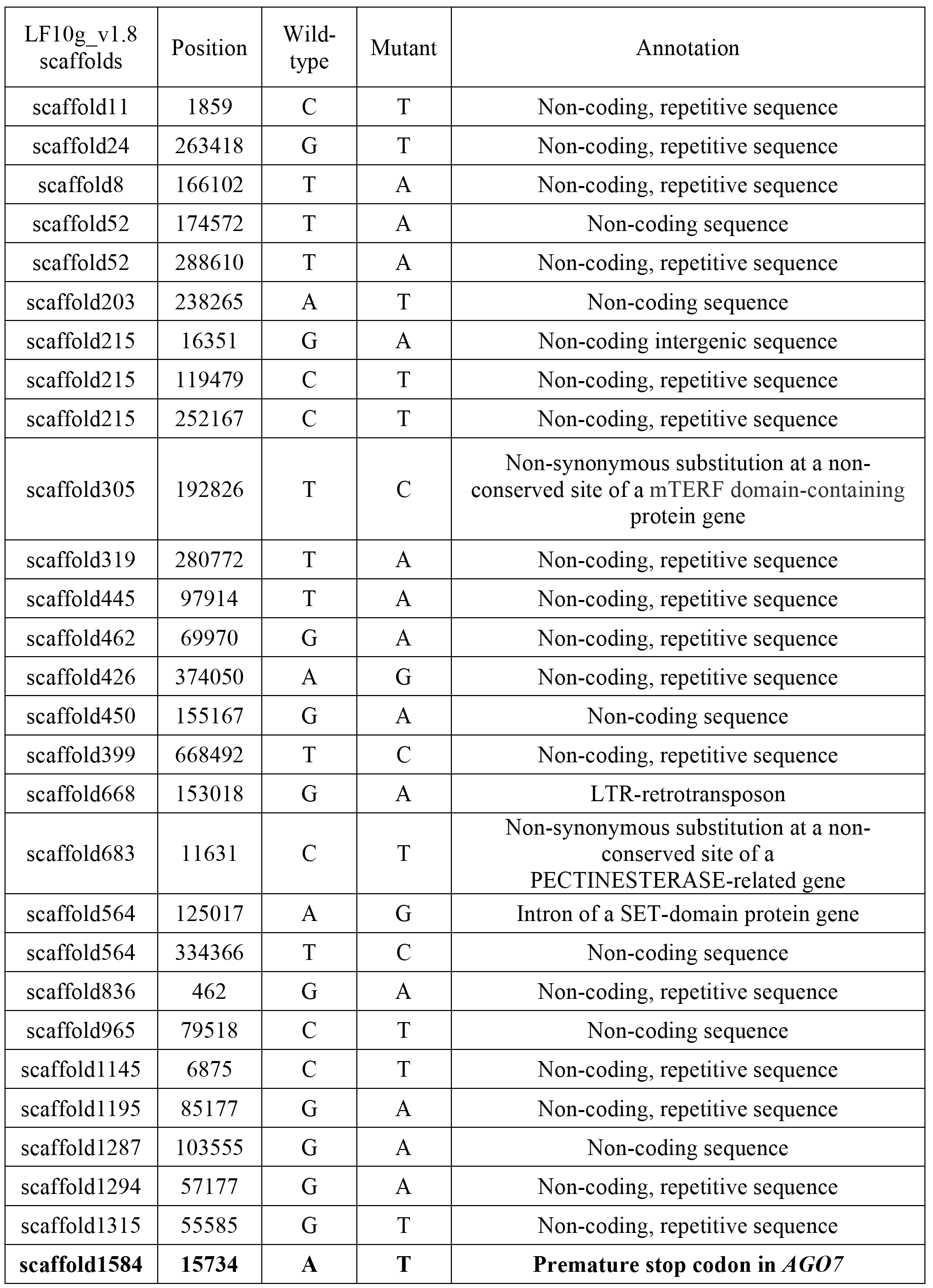
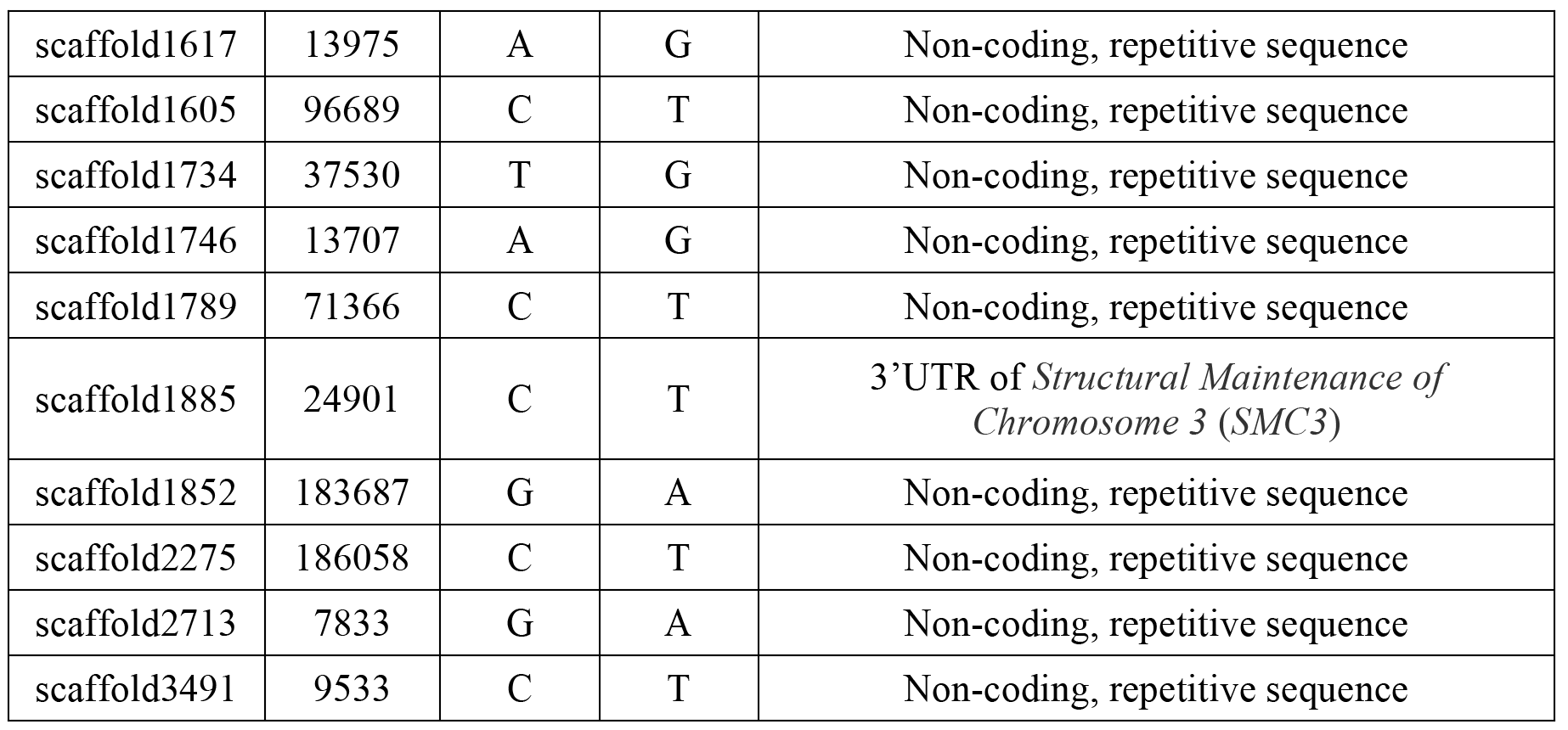
*FLAYED1* Candidate SNPs from the mutant genome comparisons. The SNP highlighted in bold is the causal mutation.

**Table S3.** *FLAYED2* Candidate SNPs from the mutant genome comparisons. The SNP highlighted in bold is the causal mutation.

| LF10g_v1.8 scaffolds | Position | Wild-type | Mutant | Annotation |
| --- | --- | --- | --- | --- |
| scaffold68 | 9337 | T | C | Non-coding, repetitive sequence |
| **scaffold156** | **87555** | **G** | **A** | **Premature stop codon in *SGS3*** |
| scaffold283 | 103550 | T | A | Non-coding, repetitive sequence |
| scaffold249 | 427385 | C | T | LTR-retrotransposon |
| scaffold432 | 106338 | G | A | Non-coding, repetitive sequence |
| scaffold432 | 202697 | A | T | Non-coding, repetitive sequence |
| scaffold427 | 215729 | G | A | Non-coding, repetitive sequence |
| scaffold925 | 11368 | C | T | Non-coding, repetitive sequence |
| scaffold1368 | 69701 | G | A | Non-coding, repetitive sequence |
| scaffold1418 | 15654 | T | A | Non-synonymous substitution at a non-conserved site of a calcium-dependent phospholipid-binding protein gene |
| scaffold1706 | 34503 | C | T | Intron of a serine/threonine protein kinase gene |
| scaffold1884 | 10588 | G | A | Non-coding, repetitive sequence |
| scaffold1977 | 48277 | T | A | Non-coding, repetitive sequence |
| scaffold1987 | 42893 | A | G | Non-coding sequence |
| scaffold1897 | 127158 | A | G | Non-coding, repetitive sequence |
| scaffold2025 | 129655 | T | A | Non-coding intergenic sequence |
| scaffold2466 | 11082 | T | A | Mutator-like transposon |
| scaffold2643 | 10512 | C | T | LTR-retrotransposon |
| scaffold2984 | 28110 | C | T | Mutator-like transposon |

**Table S4.**
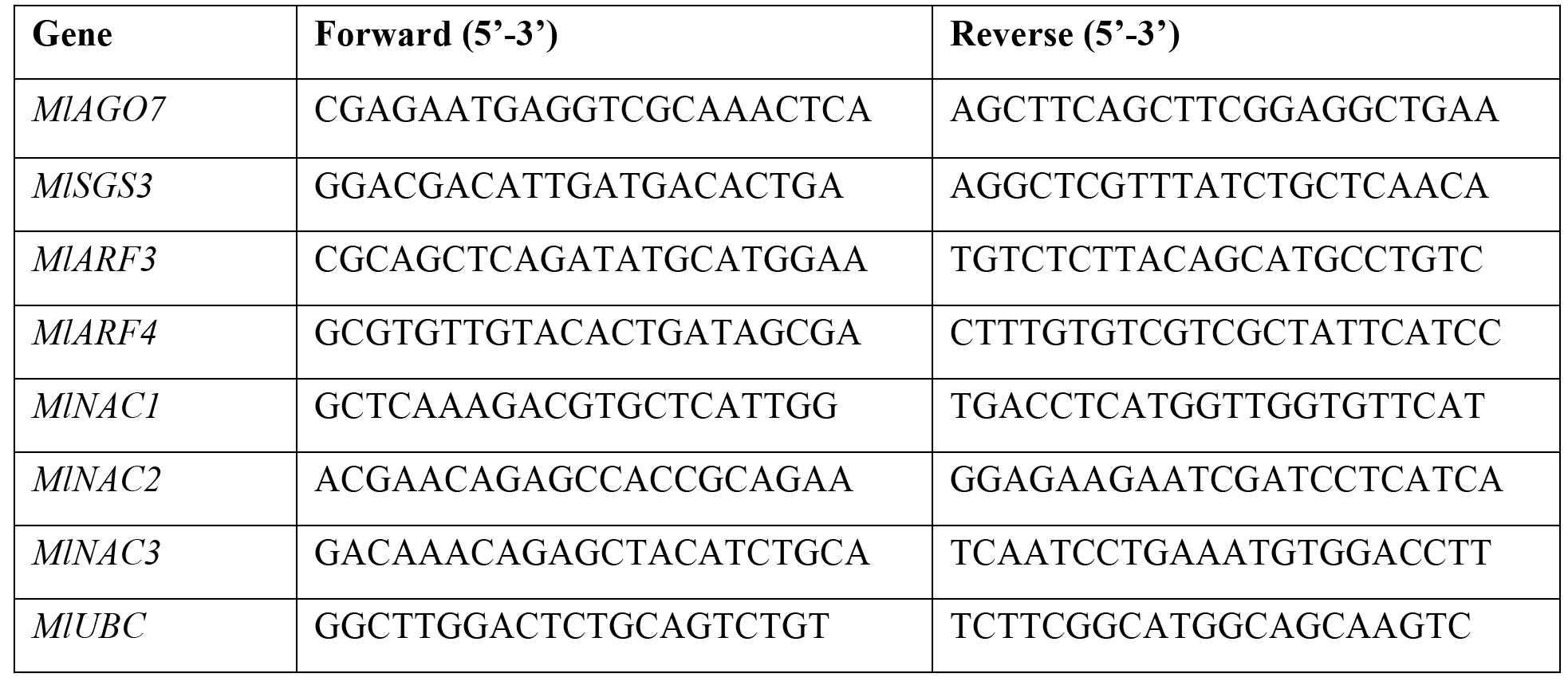
qRT-PCR Primers used in this study.

**Table S5.**
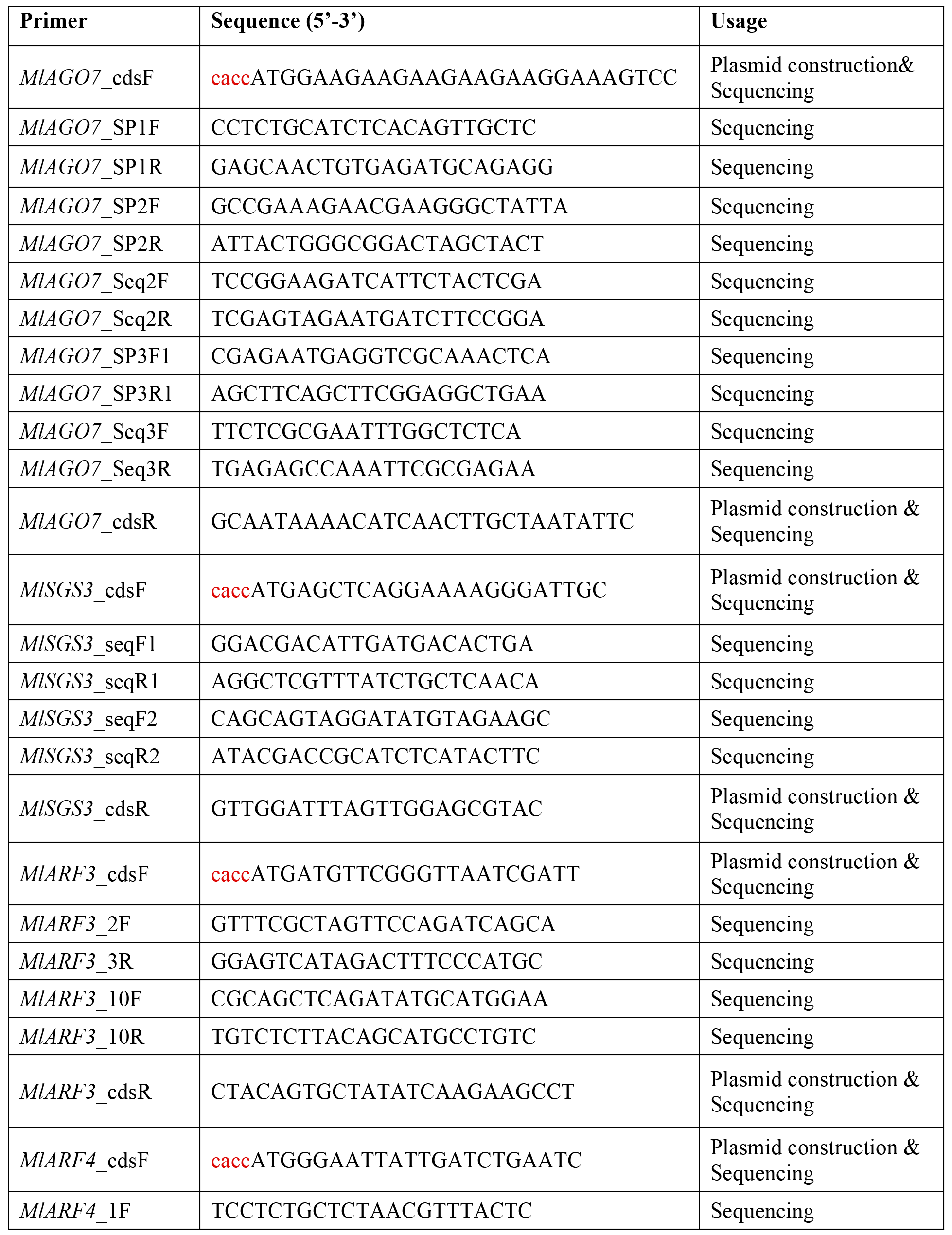
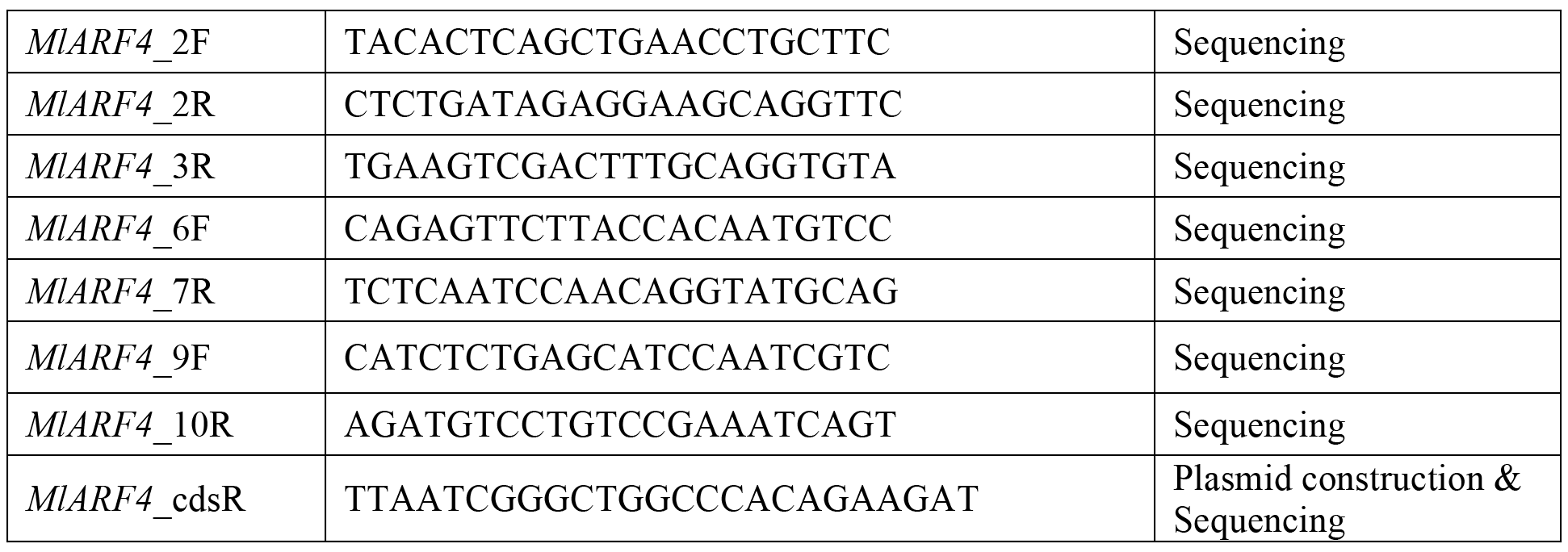
Primers used for plasmid constructions and sequence verification. The sequence highlighted in red (“cacc”) is the 4-bp sequence necessary for pENTR/D-TOPO cloning.

